# An efficient method to identify, date and describe admixture events using haplotype information

**DOI:** 10.1101/2021.08.12.455263

**Authors:** Pongsakorn Wangkumhang, Matthew Greenfield, Garrett Hellenthal

## Abstract

We present fastGLOBETROTTER, an efficient new haplotype-based technique to identify, date and describe admixture events using genome-wide autosomal data. With simulations, we demonstrate how fastGLOBETROTTER reduces computation time by 4-20 fold relative to the haplotype-based technique GLOBETROTTER without suffering loss of accuracy. We apply fastGLOBETROTTER to a cohort of >6000 Europeans from ten countries, revealing previously unreported admixture signals. In particular we infer multiple periods of admixture related to East Asian or Siberian-like sources, starting >2000 years ago, in people living in countries north of the Baltic Sea. In contrast, we infer admixture related to West Asian, North African and/or Southern European sources in populations south of the Baltic Sea, including admixture dated to ≈300-700CE, overlapping the fall of the Roman Empire, in people from Belgium, France and parts of Germany. Our new approach scales to analysing hundreds to thousands of individuals from a putatively admixed populations and hence is applicable to emerging large-scale cohorts of genetically homogeneous populations.

## Introduction

In recent years, numerous techniques have emerged that exploit expected patterns of linkage disequilibrium (LD) among Single-Nucleotide-Polymorphisms (SNPs) in admixed populations that descend from the intermixing of multiple ancestral sources, in order to identify, describe and date these admixture events. Many of these techniques assume a pulse(s) of instantaneous admixture between two or more sources, followed by random mating in the admixed population [1]. Under this model, the probability of inheriting two DNA segments from the same ancestral source along the genome of an admixed individual decays exponentially, with rate proportional to the date of admixture (in generations ago) and genetic distance between the segments [2]. This relationship is exploited by software including ROLLOFF [3, 4], ALDER [5], MALDER [6], GLOBETROTTER [2] and MOSAIC [7]. These approaches can date such admixture events, as well as estimate the proportions of DNA contributed by each admixing source. In contrast to other admixture inference techniques (e.g. [8]), an additional advantage is that they do not require accurately assigning each local segment of an admixed person’s genome to one of the admixing sources, which can be challenging in cases where admixing sources are genetically similar. These techniques and others have demonstrated admixture occuring in the last ≈4,000 years to be ubiquitous among modern human populations [5, 2].

To infer admixture, each technique uses a set of sampled reference populations that act as surrogates to the admixing sources. While ROLLOFF and ALDER identify a single best surrogate for each admixing source by finding the best model fit out of pairings of available surrogates, GLOBETROTTER and MOSAIC infer the genetic make-up of each source as a mixture of DNA from all surrogate groups, i.e. without requiring one pre-specified surrogate per source, giving these approaches more flexibility. Furthermore, while ROLLOFF, ALDER and MALDER model SNPs independently, GLOBETROTTER and MOSAIC leverage haplotype information when inferring the probabilities of descending from each admixing source for segments along an admixed individual’s genome, which can be more powerful for characterizing admixture signals when using densely genotyped or sequenced individuals [2, 9]. Also, While MALDER, MOSAIC and GLOBETROTTER can each infer multiple dates of admixture, presently among these only GLOBETROTTER can infer multiple pulses of admixture involving the same surrogate groups.

However, a key drawback of GLOBETROTTER is its computational complexity. For example, GLOBETROTTER takes ≈14 hours on a single core to identify and date admixture in a target population of 20 individuals when using 90 reference populations and ≈500,000 SNPs. Here we present *fastGLOBETROTTER*, a new method that increases the speed of inferring admixture events without sacrificing accuracy relative to GLOBETROTTER. We compare both methods using simulations of admixture events with a wide range of dates, admixture proportions and varying degrees of genetic similarity among the admixing sources. We also assess fastGLOBETROTTER’s sensitivity to demographic effects like strong bottlenecks. Finally, we apply *fastGLOBETROTTER* to a cohort of 6,209 Europeans from ten countries genotyped at 477,417 SNPs [10], inferring previously unreported admixture signals spanning Europe.

## Results

### Overview of the fastGLOBETROTTER approach

GLOBETROTTER attempts to identify, date and describe admixture in a target population of putatively admixed individuals using a set of reference populations. To do so, first the genomes of all target and reference individuals are phased using available software (e.g. [11, 12]). Next, at each SNP of the phased haploid genome *X* of a given target or reference individual, CHROMOPAINTER [13] infers which of a set of “donor” haploids shares ancestry most recently with *X*, with the most recently related donor typically the same over a string of contiguous SNPs. In this way, CHROMOPAINTER describes the two phased haploids of each target individual as a mosaic of DNA segments, with each segment matching to a single donor haploid. In practice the set of “donor” haploids is often all phased haploids from the reference populations, but they can also be only partially overlapping or entirely distinct. GLOBETROTTER then uses the CHROMOPAINTER results for all target and reference individuals to infer and date admixture in the target population, while describing the genetic make-up of each admixing source as some mixture of the reference groups.

Specifically, given the donor assignment of each DNA segment in the target individuals, GLOBETROTTER infers the probability that each segment is most recently related to each reference population, by modeling the average genome-wide CHROMOPAINTER patterns across all reference groups [2]. Then for each pair of reference populations *Y* and *Z*, GLOBETROTTER infers the probability that two DNA segments separated by genetic distance *g* have one segment most recently related to *Y* and the other segment most recently related to *Z*. After some scaling, this generates an “admixture probability curve” (also referred to as a “coancestry curve”) for *Y* and *Z* (black line in Fig 1). GLOBETROTTER jointly analyses the admixture probability curves for all pairwise combinations of reference populations, with the rate of change over *g* in all curves informative for the date of admixture, and the structure of the curves informative for the admixture proportions and genetic make-up of each of the admixing sources [2].

**Figure 1:**
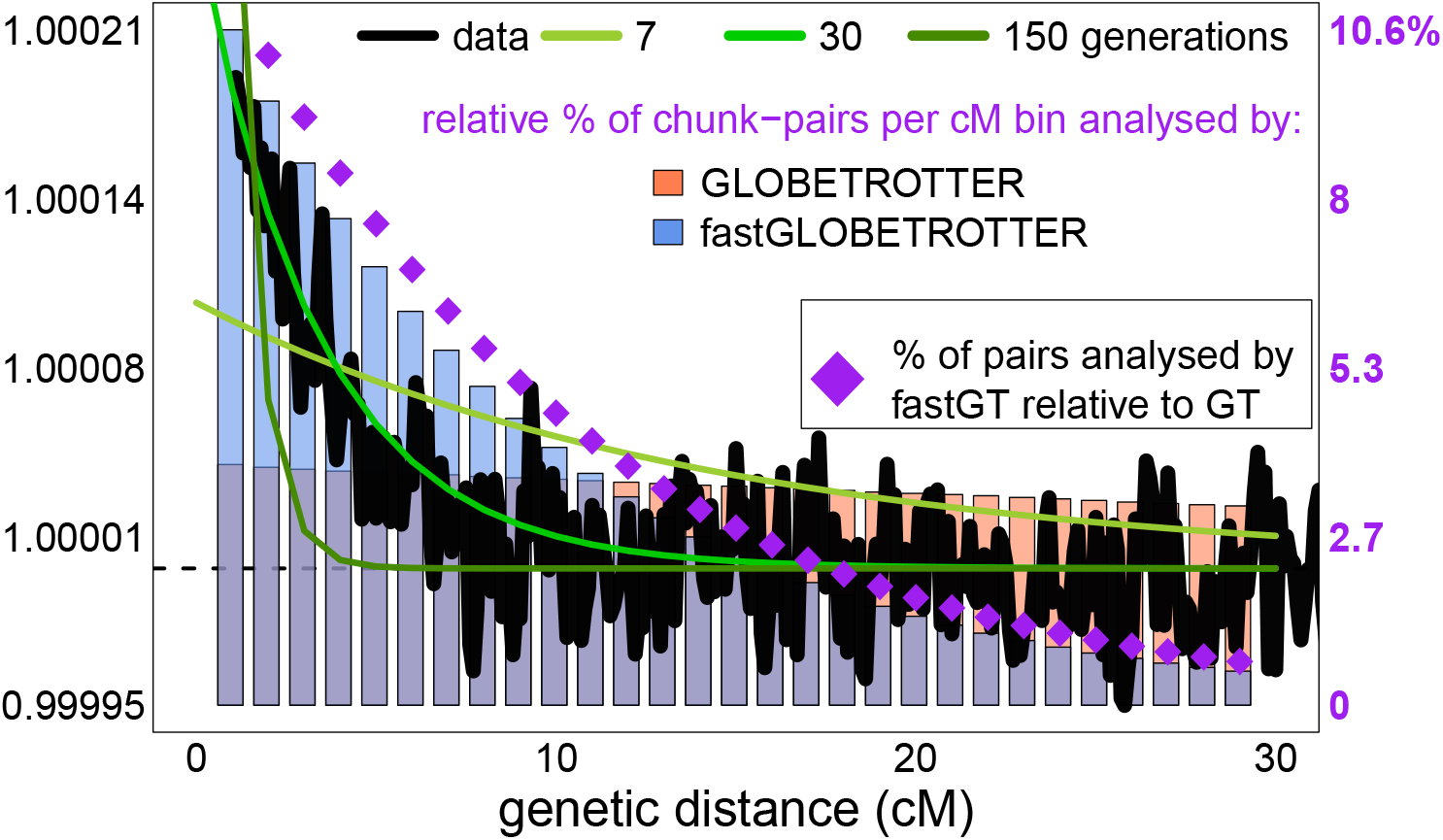
Scaled probability (black lines) that two segments separated by the given genetic distance are both inferred to share a most recent ancestor with an Irish reference individual, with key at left. These probabilities are averaged across 20 simulated individuals with European (French) and South Asian (Brahui) admixture occurring 30 generations ago. Barplots give the relative proportion of segment pairs analysed at each 1cM distance bin, in GLOBETROTTER (red) and in fastGLOBETROTTER (blue), with key at right. Purple diamonds give the proportion of segment pairs analysed by fastGLOBETROTTER relative to those analysed by GLOBETROTTER at each distance bin, with key at right. To increase computation speed, fastGLOBETROTTER analyses fewer segment pairs at each distance bin compared to GLOBETROTTER. But it preserves accuracy by analysing a higher relative proportion of the more informative segment pairs separated by smaller distances. Expected scaled probabilities for three different admixture dates are shown for comparison (green lines).

By default, GLOBETROTTER considers all pairs of DNA segments separated by (e.g.) *g <* 50cM when generating the admixture probability curve for *Y* and *Z*. However, for admixture events occurring >10 generations ago, the patterns in these curves that are attributable to admixture rapidly decline as *g* increases (green lines in Fig 1). Therefore, fastGLOBETROTTER uses a stochastic algorithm that preferentially selects DNA segments separated by short distances when generating admixture probability curves. Figure 1 illustrates this for a simulated example, with barplots comparing the relative proportions of segment pairs, separated by various distances *g*, that are included for inference under each of GLOBETROTTER and fastGLOBETROTTER. Focussing on the most informative segment pairs enables fastGLOBETROTTER to ignore other less informative pairs when constructing probability curves. We demonstrate in our simulations how this increases computational speed without sacrificing accuracy.

Our new program fastGLOBETROTTER further improves upon GLOBETROTTER in a number of additional ways. Firstly, fastGLOBETROTTER implements a technique analogous to that used in ALDER [5] to eliminate automatically signals in the admixture probability curves that are likely attributable to strong bottleneck-like effects in the target population. Secondly, fastGLOBETROTTER allows users to increase the memory used, by approximately the square of the number of donor populations, in order to further increase computational speed by approximately the number of chromosomes analysed. Thirdly, we have implemented an option to construct confidence intervals (CIs) for inferred dates by using a jackknifing approach [14], analogous to that used in ROLLOFF [4], which thus generates confidence intervals even when testing for admixture in single individuals. See Materials and Methods for more details of the fastGLOBETROTTER algorithm and features.

### Simulations show *fastGLOBETROTTER* decreases computation time without sacrificing precision

We compared the performances of the GLOBETROTTER and fastGLOBETROTTER using admixed populations from [2] consisting of 7-100 individuals simulated as mixtures of real or coalescent-simulated populations representing Africa, America, Central Asia, East Asia and Europe (Fig 2, Fig S1, Table S1, Table S2). As expected, typically performance in both is better when the admixing sources are more genetically different, when the admixture is more recent and when the fraction of ancestry from the minority contributing source is higher. The accuracy and precision is similar between GLOBETROTTER and fastGLOBETROTTER in all scenarios. Inference with fastGLOBETROTTER is equally robust to GLOBETROTTER in the presence strong bottlenecks in the target population (Table S2). The decrease in computation time for fastGLOBETROTTER depends on the number of donors and reference populations inferred to contribute to the target. Across these simulations, fastGLOBETROTTER is ≈4-12 times faster than GLOBETROTTER when inferring date point estimates and the sources and genetic make-up of each admixing population (Fig 2), if using identical memory for each approach. However, it was ≈20 times faster, with identical accuracy, if allocating more memory (in this case 1Gb) to speed up calculations (see Methods).

**Figure 2:**
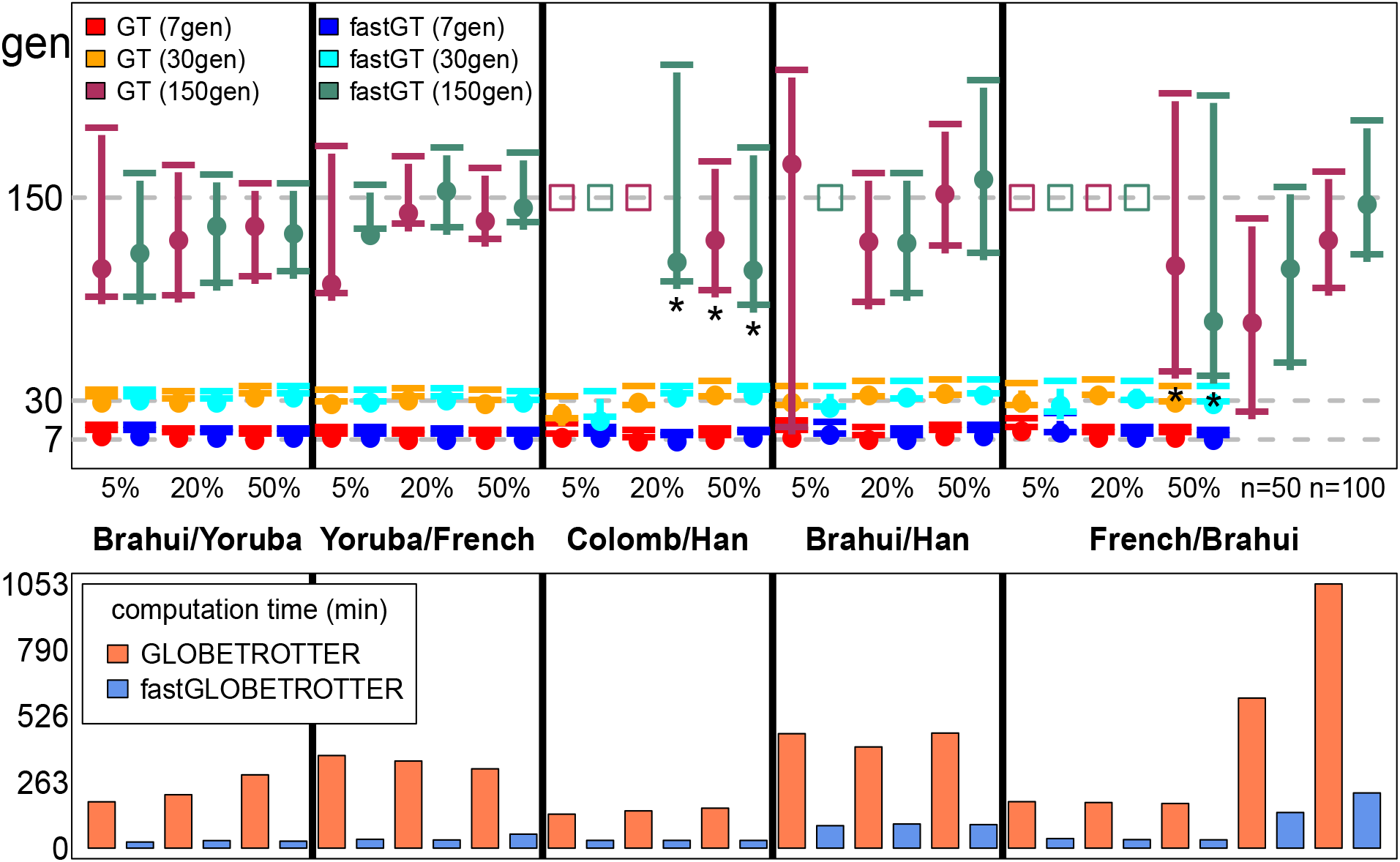
**(top:)** Inferred admixture dates (+95% CI) for simulations mixing two groups (listed in middle), with the proportion of admixture from the second group given in the x-axis. For each combination of population and admixture proportion, results are given for each of GLOBETROTTER (GT) and fastGLOBETROTTER (fastGT), for true dates of 7, 30 and 150 generations ago (grey dashed horizontal lines). French/Brahui has an additional two analyses at far right for {150 generations, 50%} when increasing the sample size *n* to 50 and 100. Cases that conclude “no admixture” are depicted with an open square placed at the true date of admixture. Asterisks (*) denote use of a different grid for binning haplotype segments (see Methods). **(bottom:)** Computation times (in minutes, excluding bootstrapping used to generate CIs) of each approach for the scenarios above, averaging across the three different admixture dates where applicable.

### Simulations illustrate limitations when inferring complicated admixture

To better understand our observed results when applying fastGLOBETROTTER to a large-scale European cohort, we also performed new simulations that mimic signals inferred in our European analysis. In these new simulations that each consider 50 admixed individuals, fastGLOBETROTTER accurately detects and describes a simple admixture event between a European source (consisting largely of people sampled from Denmark) and an East Asian (Evenk) or North African (Morocco) source occurring 100 or 200 generations ago (Table 1). However, confidence intervals around the inferred date are wide for the 200 generation old event involving North Africans.

**Table 1:** fastGLOBETROTTER inference for simulations mimicking admixture events inferred in Europeans. In each case, fastGLOBETROTTER concluded a single date of admixture between two sources. Inferred dates are given in generations. “source1” and “source2” indicate the inferred surrogate group that best genetically represents the minority and majority contributing sources, respectively, with % the inferred proportion of admixture from source 1. *R*_1_: r-squared fit of a single date (i.e. measuring fit of green line to black lines in Fig S6); *R*_2_: additional r-squared explained by adding a second date.

| simulation description | $R_1$ | $R_2$ | date (95% CI) | % | source 1 | source2 |
| --- | --- | --- | --- | --- | --- | --- |
| **Denmark 80% + Morocco 20%** |  |  |  |  |  |  |
| 100 generations ago | 0.9 | 0.03 | 106 (98-123) | 8 | ethiopiana | french |
| 200 generations ago | 0.4 | 0.05 | 157 (84-246) | 13 | egyptian | french |
| **Denmark 80% + Evenk 20%** |  |  |  |  |  |  |
| 100 generations ago | 0.94 | 0.04 | 104 (96-112) | 21 | dolgan | french |
| 200 generations ago | 0.56 | 0.08 | 221 (180-292) | 22 | dolgan | germanyaustria |
| 70 generations ago | 0.97 | 0.08 | 71 (66-76) | 25 | yukagir | germanyaustria |
| + German 20%, 10 gen ago | 0.97 | 0.23 | 61 (56-66) | 20 | yukagir | germanyaustria |

We also simulated an admixture event occurring 70 generations ago between European and East Asian sources, with and without an additional pulse of admixture 10 generations ago involving another European source (primarily consisting of people sampled from Germany) (Fig S2). However, fastGLOBETROTTER only detects a single admixture event in the case of two simulated pulses of admixture. This is not surprising, as two admixture pulses that occur relatively close in time are difficult in theory to disentangle from a single admixture event with a date between these two pulses or continous admixture between the same sources [2]. However, the fit to the data when assuming two pulses of admixture increases by ≈3-fold when we have simulated two pulses relative to only one pulse (*R*_2_ in Table 1), which provides a potentially useful indicator of more than one admixture pulse. In addition, in the case of an additional pulse recent admixture, the inferred date is more recent (61 generations ago; 95% CI: 56-66) relative to when there is no additional recent admixture (71 generations ago; 95% CI: 66-76). This suggests that, in the case of two pulses of admixture, the inferred event assuming one pulse lies somewhere between the initial and most recent admixture event.

### Admixture events in Europe spanning 50BCE-1400CE

We applied fastGLOBETROTTER to 6,209 Europeans from ten countries described in [10, 15]. To account for putative structure within this cohort, we first used fineSTRUCTURE [13] to cluster these Europeans into 86 genetically homogeneous groups, which ranged in sample size from 9 to 212 individuals (Table S3). We then applied fastGLOBETROTTER to each cluster separately, using 162 reference populations as potential surrogates for putative admixture events [10, 16, 2, 17, 18, 19, 20, 21, 22, 23, 24, 25] (Table S4). FastGLOBETROTTER detected admixture in 80 of the 86 clusters, inferring a simple event between two sources at a single date in 60 clusters, >2 sources admixing at around the same time in 17 clusters, and multiple dates of admixture in three clusters (Table S5). Inferred dates range from 18 to 64 generations ago, ≈600-2000 years ago assuming 28 years per generation [26].

Inferred sources of ancestry can be categorised broadly into two classes that are highly correlated with geography (Fig 3, Fig S3, Fig S4, Fig S6-S16). The first involves ancestry related to West/Central Asia, North Africa and/or Sub Saharan Africa, which is found to varying degrees (3-36%), along with southern European-like ancestry, in nearly all clusters predominantly comprised of people from Belgium, Denmark, France, Germany, Spain and Italy. The second is ancestry related to East Asians and Siberians, of which 3-17% ancestry is found in nearly all clusters containing people from Finland, Norway and Sweden, plus the cluster (C37) containing the majority of sampled Polish individuals.

**Figure 3:**
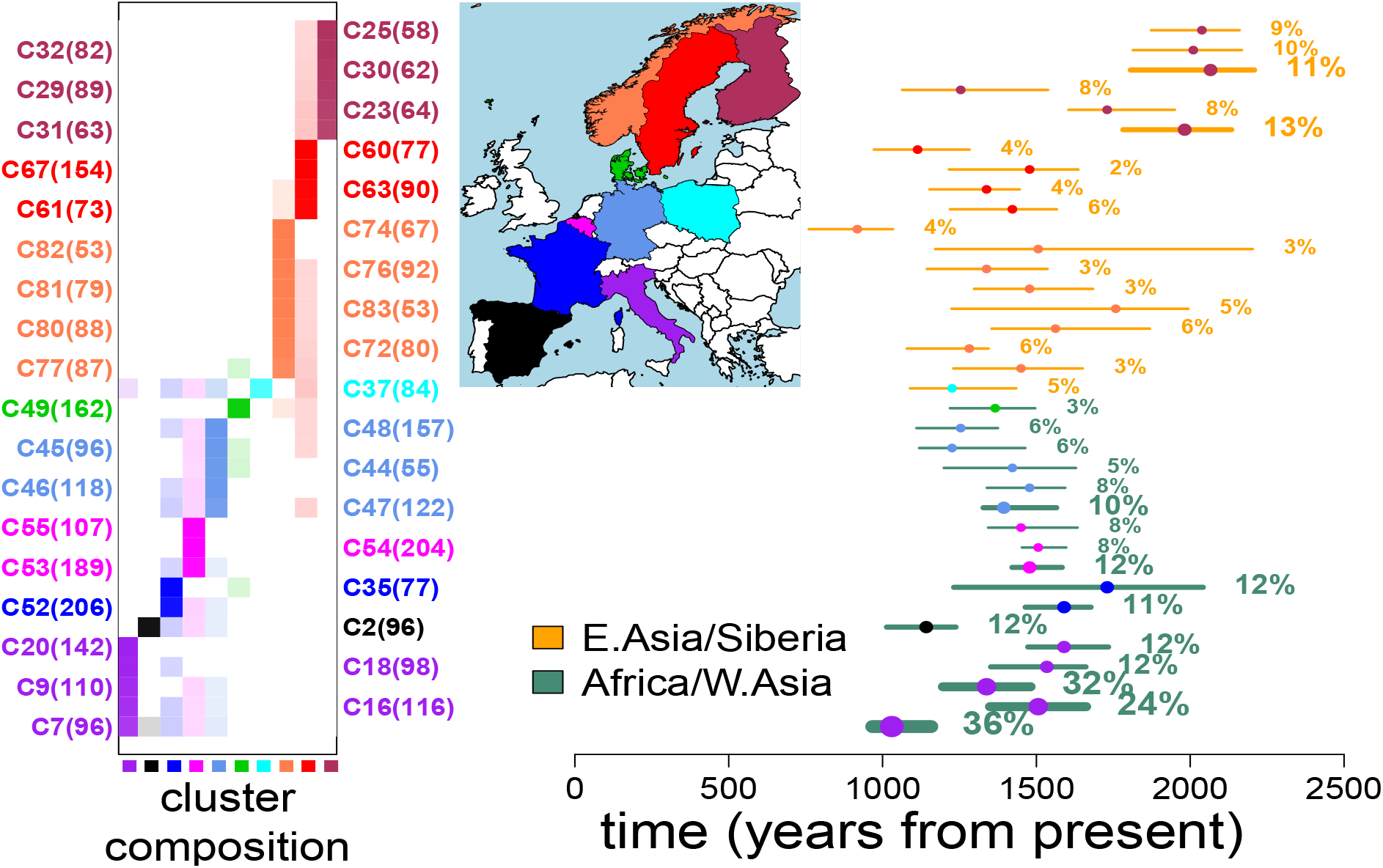
**(right:)** Inferred admixture dates (+95% CIs) for the 36 Europe clusters that contain >50 individuals, have a single inferred date, and have a >2% inferred contribution from reference groups in E.Asia/Siberia (orange CIs) or in Africa/W.Asia (teal CIs). The inferred contributions from these two regions are given to the right of each CI; *“Africa/W*.*Asia”*={West Africa, West Asia, South Middle East, North Africa, San, Central Africa, East Africa, Bantu, Ethiopian} and *“E*.*Asia/Siberia”*={Siberia, Northeast Asia, Southeast Asia} as defined in Table S4. **(left:)** The proportion of individuals in each cluster (row) from each country (column), with clusters’ sample sizes in parentheses and countries’ colors in the map. Each cluster’s label at left and point estimate date at right is colored according to the country most represented in that cluster.

Thirty-six clusters had >50 individuals, a single inferred admixture date, and >2% inferred ancestry from the above sources (Fig 3). Among these 36, the highest degrees (> 20%) of inferred ancestry related to West Asians and North/Sub-Saharan Africans are found in three clusters that predominantly consist of Italians, with inferred dates covering a broad range spanning ≈1000-1500 years ago (Fig 3). Interestingly, seven of these 36 clusters, consisting of Belgians, French, and Germans (clusters C44, C46, C47, C52-C55), have similar inferred amounts (5-12%) of ancestry related to North African and West Asian (Armenian, Morocco, Turkey) sources and similar admixture dates of ≈300-700CE (Fig 3, Fig S4).

In contrast, the 18 of these 36 clusters primarily consisting of individuals from countries north of the Baltic (though with cluster C77 containing three Danish individuals) have inferred admixture events dated to ≈900-2100 years ago involving a source related predominantly to present-day Russians and Siberians. Of these 18 clusters, inferred dates are oldest in four of the predominantly Finnish clusters (C25, C30-C32), and significantly more recent in nine of the twelve clusters primarily containing Swedes and Norwegians. Furthermore, 12 of these 18 clusters have some evidence of multiple pulses of admixture (i.e. *R*_2_ > 0.2 in Table S5).

## Discussion

Here we introduce fastGLOBETROTTER, a new program to identify, date and describe admixture events with 4-20 times increased computational efficiency over GLOBETROTTER. Our simulation results suggest that this computational increase comes without a loss of accuracy and precision. Indeed we see evidence of fastGLOBETROTTER outperforming GLOBETROTTER in some of the more challenging scenarios, such as when simulating admixture occurring 150 generations ago between the relatively genetically similar French and Brahui. In this scenario, fastGLOBETROTTER’s point estimates are closer to the truth relative to GLOBETROTTER’s when simulating 50 or 100 admixed individuals (Fig 2). In principle, fastGLOBETROTTER’s sub-sampling scheme, which down-weights pairs of distantly-separated DNA segments when constructing the admixture probability curves, could increase accuracy over GLOBETROTTER’s approach that does not downweight such segments, particularly for older events. This is because distantly-separated segments provide little to no information about the admixture event (Fig 1), while being more susceptible to random noise due to the smaller number of segments separated by such large distances.

The 86 fineSTRUCTURE-inferred clusters of 6,209 Europeans were largely consistent with country label (Table S3) and likely reflect the high degree of geographic clustering previously observed in these data [15] and using other data from these countries [27, 28, 29, 30, 31, 32, 33, 34]. While we do not have access to any fine-scale geographic information beyond the country level, we observe broad geographic trends in inferred ancestry across these European clusters. In particular clusters nearly exclusively containing people from countries north of the Baltic Sea (i.e. Finland, Norway, Sweden) exhibit 2-13% inferred ancestry from a source most closely related to East Asians and Siberians. In contrast, clusters containing primarily individuals from south of the Baltic (i.e. Belgium, Denmark, France, Germany, Italy, Spain), except the cluster containing the majority of Polish individuals, have *<*1% inferred ancestry related to East Asians and Siberians, instead exhibiting 3-36% inferred ancestry from sources related to Central/West Asia, North Africa and/or Sub-Saharan Africa, along with inferred ancestry related to southern Europe (Fig 3, Fig S4, Table S5).

The signals in Finland are consistent with Finns descending from an early intermixing between European and E.Asian/Siberian-like sources. Plausibly related intermixing was previously reported by Saag et al., 2019 [35], who found evidence of Siberian-like admixture appearing in ancient DNA (aDNA) samples from nearby Estonia in Late Bronze Age graves ≈2,500 years ago. While their reported date is older than that inferred here by fastGLOBETROTTER, our results do not preclude multiple episodes of intermixing involving an East Asian-like source present in the region by the Iron Age. In particular our simulations illustrate how multiple dates of admixture between two sources, where a group genetically similar to one of the original admixing sources subsequently intermixes with the previously admixed group, may be inaccurately described by fastGLOBETROTTER as a single admixture event, sometimes with an inferred date somewhere between the dates of the two admixture events (Figure S2, Table 1). Supporting this, each of the four Finnish clusters with mean inferred date older than 50CE (C25, C30-C32) show some evidence of multiple pulses of admixture (*R*_2_ > 0.2 in Table S5) consistent with what we see in simulations with multiple admixture pulses (Table 1).

When moving geographically from Finland to Sweden and Norway, i.e. east to west, clusters of people north of the Baltic show more recent inferred dates and decreasing proportions of inferred ancestry related to E.Asia/Siberia (Fig 3). These observations are consistent with a scenario where a source related to northwest Europeans initially intermixed with a source related to East Asians around (or plausibly older than) 50BCE, with this initial intermixing occurring geographically nearer to Finland than to Norway or Sweden. Subsequently, this admixed group could have intermixed with other unadmixed Europeans through migrations westward, a process that could lead to the decreased date estimates and decreased E.Asian/Siberian proportions of ancestry we infer in Norway and Sweden (Fig 3) mimicked by our simulations (Figure S2, Table 1). Larger sample sizes from these areas, which may allow fastGLOBETROTTER to correctly identify and date multiple pulses of admixture, and/or additional data from ancient human remains may shed light on whether this is indeed the case.

The wide range of inferred dates and ancestry proportions in Italy is consistent with multiple episodes of intermixing with sources related to present-day peoples from West/Central Asia, North Africa and/or Sub-Saharan Africa as has been previously reported [2, 16, 32]. In the one cluster (C2) consisting primarily of Spanish in Figure 3, the inferred date of ≈1200 years ago involving sources with Sub-Saharan African DNA matches previous reports of intermixing potentially related to the Muslim conquest of Spain [31].

The majority of clusters consisting of Belgians, French, and Germans (C43-C48, C51-C55) are inferred to have one-date of admixture between two sources, with one genetically related to the modern-day British and Norwegians, and the other genetically related to modern-day southern Europeans (Greeks, Italians), Cypriots, Moroccans and/or Armenians (Figure S3, Figure S4, Table S5). The inferred point estimate dates of these events span a relatively small range of 400-650CE (Table S5), though slightly later (750-800CE) in clusters C45 and C48. While the historical events driving these signals are unclear, a plausible explanation is that it relates to the Roman Empire, which covered all of present-day Belgium, Germany, France, Turkey, North Africa and elsewhere prior to its decline and eventual fall in 476CE [36]. In particular individuals carrying ancestry recently related to that found in present-day people from North Africa, West Asia and southern Europe could have moved across the empire during this time. A recent study of aDNA samples found in or near Rome spanning the time of the empire (27BCE-300CE) reported signals of ancestry from disparate sources genetically related to present-day people from the near East, eastern Mediterranean and North Africa [37], suggesting the Empire faciliated migrants into Rome during this time. Our results, based on analysing genetic variation data from present-day individuals, are consistent with this migration extending across the empire, with individuals carrying such ancestry intermixing with people living in or around present-day Belgium, France and Germany either during or soon after the fall of the Roman Empire.

Our simulations suggest our inferred dates are biased to detect recent admixture (Table 1), indicating admixture we detect in these northwest Europeans may have begun prior to our inferred dates of ≈400-650CE. However, a previous study reported genetic patterns in aDNA from Bavarians dated to ≈500CE showed no clear genetic affinities to present-day southern Europeans, consistent with a lack of widespread intermixing between local and southern sources in Germany prior to our inferred admixture dates [38]. This was despite that study reporting the presence of a presumed Roman soldier in Munich dated to ≈300CE, which had strong genetic affinities to present-day southern Europeans [38]. Additional aDNA samples from these northwest European regions may help clarify these signals.

Other German clusters (C38, C39, C41, C42) do not appear to have this S.Europe/W.Asian/African signal, instead exhibiting inferred signals of ancestry sources related to eastern European groups such as Poland (Table S5) and significantly more recent dates around 1100-1400CE, perhaps reflecting geography.

The computational speed-ups described here are a step change over GLOBETROTTER. For example, when using its fastest option on a single computing node, fastGLOBETROTTER took just under 21 hours, using ≈9Gb of RAM, to perform date/source inference and 100 bootstrap resamples using default values on European cluster C9 containing 110 individuals. In contrast, using the same input parameters, GLOBETROTTER completed the date/source inference step and only 30 bootstrap re-samples after 30 days. These speed-ups enable a far more efficient analyses of larger datasets, and hence detection of more subtle admixture signals. Nonetheless, future improvements are necessary to scale to e.g. many thousands of individuals. Furthermore, the computational gains described here apply only to the GLOBETROTTER inference step, and not to the phasing and chromosome painting steps prior to applying GLOBETROTTER. However, we note that phasing algorithms are relatively fast, with current software able to phase tens to hundreds of thousands of individuals in a few hours on a computational cluster. CHROMOPAINTER typically is computationally much slower than phasing, but is parallelisable by both target individual and chromosome in a manner GLOBETROTTER and fastGLOBETROTTER are not, with alternative “chromosome painting” software existing that is considerably faster than CHROMOPAINTER (e.g. [39]). Given the increasing ubiquity of large-scale genotype resources from relatively genetically homogenous populations (e.g. [40, 41]), such computational speed-ups will become increasingly necessary to cope with present-day sample collections.

## Materials and Methods

### Details of DNA segment pair selection algorithm in fastGLOBETROTTER

The theory behind fastGLOBETROTTER is analogous to that in GLOBETROTTER, which has been previously described [2]. In brief, consider an admixed population that descends from the mixture of two source groups *A* and *B* that contributed *α* and 1 − *α* of the DNA, respectively, and intermixed *λ* generations ago. Assuming random mating among admixed individuals since the time of admixture, and that that the crossovers between any two loci occur at random (i.e. no crossover interference) [1], the probability *P*_*A→B*_(*g* | *λ, α*) that two loci separated by genetic distance *g* (in Morgans) along a chromosome within a haploid genome of an admixed individual, with one locus descending from an individual from *A* and the from an individual in *B*, is:

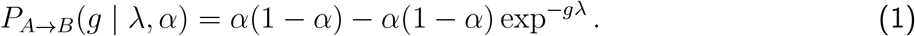

The probability *P*_*A→A*_(*g* | *λ, α*) that the two loci both descend from individuals from *A* is:

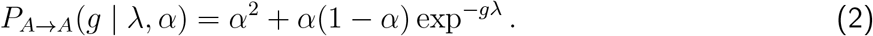

(Analogous formulas can be derived for more than two admixing sources [2].) Note that *α*^2^ is the marginal probability that two independent loci (e.g. separated by a large distance *g*) both derive from source *A*, with *α*(1 − *α*) similarly the marginal probability that two independent segments derive from *A* and *B*. Dividing (1) and (2) by *α*(1 − *α*) and *α*^2^, respectively, gives:

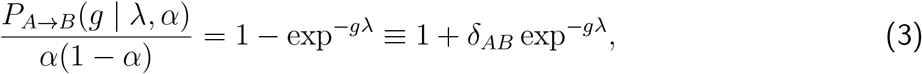

and

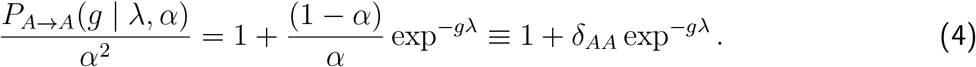

Importantly, *δ*_*AB*_ *<* 0 and *δ*_*AA*_ > 0, ensuring that equation (3) is monotonically increasing with *g* while (4) is monotonically decreasing with *g*. While the true admixing sources *A* and *B* are unknown, we instead consider the probability that two segments separated by distance *g* are inferred to share most recent ancestry with reference populations *U* and *V*. If *U* and *V* are good surrogates for the same source (e.g. source *A*), then this scaled probability curve will be decreasing. In contrast, if *U* is a good surrogate for source *A* and *V* a good surrogate for *B*, this curve will be increasing. GLOBETROTTER and fastGLOBETROTTER exploit these relationships to infer *α* and the genetic make-up of each source group as mixtures of the reference populations, with the shape of the scaled probability curves used to infer *λ*, as described in [2].

An example of (4) is given for simulations in Fig 1, with examples of (3) for real data given in Figures S6-S16. We refer to these as “admixture probability curves”. We use CHROMOPAINTER [13] to compose each phased haploid of individuals from a putatively admixed population as a sequence of non-overlapping DNA segments, where each segment is inferred to share most recent ancestry with a reference population. Technically, a DNA segment is defined as a contiguous set of SNPs in the target haploid that are inferred by CHROMOPAINTER to share most recent ancestor with the same donor haploid. In practice, CHROMOPAINTER generates *s* (typically *s* = 10) inferred genome-wide sequences of these segments for each target haploid, giving 2*s* inferred sequences across an individuals’ two haploid genomes. Assume that these 2*s* sequences contain *C* DNA segments in total for the individual.

To construct admixture probability curves capturing (3) and (4), GLOBETROTTER uses all 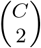 pairings of DNA segments within an individual that are (i) on the same chromosome and (ii) separated by *K* ≤ centimorgans (e.g. *K* = 30). This is among the most computationally intensive steps of GLOBETROTTER, with complexity squared in the maximum number of DNA segments matched to the same reference population across chromosomes. In contrast, fastGLOBETROTTER only uses a sub-set of all possible pairings of the *C* DNA segments.

The sub-sampling algorithm of fastGLOBETROTTER is as follows:

1. Divide a the haploid genome of an admixed individual into *B* non-overlapping bins of size *X*cM. *X* is the *bin*.*width* specified by the user; here we use *X* = 0.1 unless otherwise noted.
2. Find which of the *C* total segments fall into bin *G*_*i*_ for all *i* ∈ [1, …, *B*]. A segment will be put into bin *G*_*i*_ if the midpoint of the segment falls within the range of bin *G*_*i*_. Let *N*_*i*_ be the number of segments within bin *G*_*i*_, where 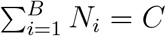.
3. For each bin *G*_*i*_, the program will compare the segments in this bin to those in bin *G*_*i*+1_, where the distance *D*_*i→i*+1_ between *G*_*i*_ and *G*_*i*+1_ is *X*cM. The program then compares *G*_*i*_ with *G*_*i*+2_ (i.e. with distance *D*_*i→i*+2_ = 2*X*) between them) and so on, until reaching the last bin *n* with *D*_*i→n*_ ≤ *K*, where *K* is the maximum allowed distance between segments. Here we use *K* = 30cM, noting that segments on different chromosomes are never compared. Also note that segments within the same bin *G*_*i*_, which by definition have distance between them *< X*cM, will not be compared to each other. However, this seems desirable as segments at such short distances can be confounded by background linkage disequilibrium unrelated to the admixture signal. The user can also specify to avoid fitting segments separated by less than some distance; here we use the default value of not fitting segments separated by ≤1cM.
4. To do the comparison in step 3, we do the following: 4a. For each *i* and *j*, where *i < j*, calculate *Y*_*ij*_ = *N*_*i*_ * *N*_*j*_ * *M*_*ij*_, which is the number of samplings of segment pairs from bins *G*_*i*_ and *G*_*j*_ to be performed, i.e. with one segment sampled from bin *G*_*i*_ and the other sampled from bin *G*_*j*_. *M*_*ij*_ is a scalar that is derived from a distribution that allows us to sample a different proportion of the total segment pairs in bins *G*_*i*_ and *G*_*j*_. For example, if *M*_*ij*_ = 1, a roughly equivalent number of segment pairs will be sampled as in the original GLOBETROTTER. Alternatively, one could make *M*_*ij*_ *<* 1 and set a higher value of *M*_*ij*_ for segment pairs with smaller *D*_*i→j*_, meaning closer pairs are preferentially sampled over more distant pairs. Here we use 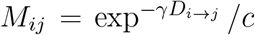, with *γ* = 0.05 and *c* = 8, which performs well in simulations in terms of computation decrease while maintaining precision (Fig 2). With these values, the number of segments pairs sampled is ∼6.5% of the total possible pairs separated by ≤30cM. Figure 1 depicts *γ* = 0.1 and *c* = 8. 4b. To compare segments in *G*_*i*_ and *G*_*j*_, the program randomly samples *Y*_*ij*_ segment pairs without replacement, with one segment from *G*_*i*_ and the other from *G*_*j*_.
5. Repeat step 4 for all pairs of bins (*G*_*i*_, *G*_*j*_) across the chromosome separated by ≤ *K*cM.

In addition, Hellenthal et al 2014 describe a “null individual” analysis that aims to eliminate LD decay signals in the admixture probability curves that are not attributable to admixture, hence providing more reliable date estimates [2]. This is done by building a “null” probability curve using segment pairs where each segment is from the painting sample of a different target individual. This “null” probability curve should be unrelated to the admixture event, because segment pairs on different individuals cannot fall within a single block of DNA inherited intact from an admixing source individual. GLOBETROTTER scales each admixture probability curve by this “null” probability curve before inferring dates and proportions of admixture, which can lead to more accurate inference, particularly in cases where the target population has experienced a strong bottleneck [2] (e.g. see Table S2). To implement this “null” individual protocol into fastGLOBETROTTER, we replace the following two steps in the above algorithm:

- Step 2. For *T* admixed target individuals, let *P*_*null*_ be a vector of size 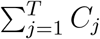, with *C*_*j*_ the total number of segments for target individual *j*, and where *P*_*null*_(*c*) stores the index of the admixed target individual to which segment *c* belongs.
- Step 4b. To compare segments in *G*_*i*_ and *G*_*j*_, the program randomly samples segment pairs, with one from *G*_*i*_ (call this segment *a*_*i*_) and the other from *G*_*j*_ (call this segment *b*_*j*_). When building the “null” probability curve, we only consider segment pairs where *P*_*null*_(*a*_*i*_) ≠ *P*_*null*_(*b*_*j*_), with segments randomly chosen until *Y*_*ij*_ pairs meet this criterion.

After constructing the “admixture probability curves” as described above, the inference steps in fastGLOBETROTTER are the same as those described in [2] for GLOBETROTTER, except the few changes detailed below.

### Details of approach to adjust for extensive linkage disequilibrium within the target population in fastGLOBETROTTER

In admixed groups affected by strong bottlenecks, DNA segments matched to donors using CHROMOPAINTER may be atypically long, reflecting high levels of within-population linkage disequilibrium (LD). To cope with this, GLOBETROTTER ignores any chunk pairs separated by ≤1cM when generating admixture probability curves, implicitly assuming such within-population LD is unlikely to extend beyond 1cM. However, this threshold is arbitrary; other methods like ALDER [5] attempt to automatically identify the threshold of minimal distance between segments to use when capturing admixture linkage disequilibrium. A particular concern is that the presence of many atypically long chunks over 1cM can mimic patterns in the admixture probability curves that are similar to those expected under multiple distinct pulses of admixture involving different groups admixing at different times (see [2]), hence leading to inaccurate admixture inference.

To cope with this issue, prior to model fitting, fastGLOBETROTTER automatically removes the left-end portions of the admixture probability curves that are believed to be affected by within-population LD. To do so, we first analyse the admixture probability curve of the surrogate group inferred to contribute the highest proportion of ancestry, with is likely be the most informative and clear curve. This curve provides the scaled probability that two segments separated by distance *g* both match to this surrogate group, with *g* binned to the nearest (e.g.) 0.1cM. In the absense of within-population LD, this curve should be monotonically decreasing. Therefore, starting at the left-most distance grid-point *x*, we fit a simple linear regression of the scaled probability versus distance for a window of *M* adjacent distance values, i.e. fitting a linear regression from distance grid-points *x* to *x* + *M*. The value of *x* is user-supplied; in all analyses here we use the grid-point corresponding to 1cM, meaning that two segments separated by *<* 1cM are ignored. If the fitted slope of this regression is ≥0, we shift the window one distance bin to the right, i.e. now fitting a linear regression from distance grid-points *x* +1 to *x* + *M* +1. We repeat this process until the fitted slope is *<* 0. Letting *x*_*l*_ be the left-most distance grid-point in this *M*-length window where the regression slope was *<* 0 for the first time, we record *x*_*m*_ = *x*_*l*_ + ceiling(*M/*2) as the right endpoint of the region to remove. We repeat this protocol for windows of size *M* = {3, 5, 7, 9, 11, 13}, and use the maximum value of *x*_*m*_ across all *M* as the left-end portion we eliminate from all admixture probability curves prior to inferring admixture. (In its current implementation, at most half of the total fitted distance specified by the user can be removed.) We note that considering only the admixture probability curve of the maximally-contributing surrogate group may not be sufficient to adjust for within-population LD effects in all admixture probability curves, though this approach worked well in practice. In this paper, we used this default LD-removal step for the analyses of all European populations and the simulated European populations.

### Details of memory and speed trade-off in fastGLOBETROTTER

One of the steps of GLOBETROTTER, unaffected by the speed-ups mentioned above, has computational cost that is squared in the number of donor groups included in the CHROMOPAINTER analysis. This calculation is done once per individual per chromosome in GLOBETROTTER. In contrast, fastGLOBETROTTER gives the option of performing this calculation once per individual for all chromosomes, which in practice we found to be ≈2 times faster than the standard fast-GLOBETROTTER while giving the same results. However, this approach incurs a memory increase equal to the square in the number of donor groups divided by the square of the number of surrogate groups that contribute >0.5% of ancestry. We have implemented an option in fastGLOBETROTTER which does a quick estimate of the amount of memory increase necessary to perform this new step, enabling users to decide whether the speed-up is worth the memory increase.

Due to the above, reducing the number of donor groups used by fastGLOBETROTTER can alleviate both the computational and memory constraints. Here we have also implemented an option to combine donor groups that share a similar genetic background. To do so, we first find the average proportion of genome-wide DNA that individuals from each surrogate group match to haploids from each donor group, standardising this to sum to 1.0 in each surrogate. Thus we describe each surrogate by a vector of length equal to the number of donor groups. Conversely, each donor group can be defined by a vector containing the amount they contribute to each surrogate group. For all pairs of donors groups, we find the Pearson correlation of these donor vectors. For donor vectors with correlation > 0.95, we merged their values within each surrogate group by averaging those donors’ contributions to that surrogate group. However, for the applications considered here, we found little additional reduction in computation time or memory when using this approach relative to that described in the previous paragraph, cautioning that it may not be worth the potential loss in power from joining donor groups.

### Details of jack-knifing procedure to infer confidence intervals for dates in fastGLOBETROTTER

The algorithm GLOBETROTTER uses bootstrap re-sampling of individuals to construct confidence intervals for inferred admixture dates. However, bootstrap re-sampling is not possible when inferring admixture in a single individual. Therefore, follwing [4] fastGLOBETROTTER also includes an alternative jackknifing procedure that instead drops one chromosome at a time and estimates the dates using data from the other 21 chromosomes. This process provides 22 estimated date values, which can then be used to give confidence intervals for the inferred admixture date using previously derived jack-knifing formulas (e.g. [14]). We provide a comparison of standard deviations calculated from bootstrapping versus jackknifing in Figure S5 for the European clusters. The values are notably correlated, though jack-knifing gives larger values overall as expected [42], suggesting this should only be used when testing for admixture in populations with small samples sizes.

### Simulations from Hellenthal et al 2014 [2]

To compare fastGLOBETROTTER to GLOBETROTTER, we used two sets of simulated datasets described in [2]. In the first case, each simulated haploid genome of the admixed individuals was generated as a mosaic of DNA segments from the phased genomes of individuals from the two admixing source populations, using the technique in [43] and real sampled individuals for the source populations. The following combinations of populations were used as the two admixing sources: Yoruba (Africa) versus French (Europe), French versus Brahui (West Asia), Brahui versus Han (East Asia), Brahui versus Yoruba, and Colombian (America) versus Han. For each of these population pairs, 7 (Colombia vs Han) or 20 (all others) individuals were simulated for each combination of admixture date at 7, 30 and 150 generations ago and fraction of admixture from the second source of 0.05, 0.2 and 0.5 (Fig 2, Table S1). In particular, the size of each DNA segment for an admixed haploid was sampled randomly from an exponential distribution with rate equal to the date of admixture, and this segment was copied intact from a source haploid chosen randomly from an admixing source population with probability equal to the desired admixture fraction from that source population. Here we also created new simulations of 100 admixed individuals in this manner for the case of admixture between French and Brahui at 150 generations ago and an admixture fraction of 0.5. In each case, we inferred admixture using the 91 other sampled populations from [2] as reference populations and using the CHROMOPAINTER procedure described in that paper.

The other set of simulations from [2] first used the program macs [44] to simulate 11 populations meant to mimic the demographic history of Africa (Pops 1-4), West Eurasia (Pops 5-7) and East Asia (Pops 8-11) (see Fig S1). Next, an admixed population was generated by randomly sampling 100 haploid genomes from a population comprised of 150 and 100 simulated haploid genomes from populations 2 and 8, respectively. Thus this admixed population mimics a scenario with 60% and 40% ancestry inherited from Africa and East Asia, respectively, with its small population size (50 individuals) mimicking the effects of a strong bottleneck. This admixed population was then simulated forwards-in-time, by randomly selecting parents in the current generation to construct each offspring’s two haploid genomes in the next generation, mixing the parental genomes according to recombination probabilities from the HapMap Phase 2 genetic map and generating 50 offspring in total. Separate simulations ran this forwards-in-time procedure for 10, 20 and 45 non-overlapping generations, mimicking different dates of admixture. To infer the admixture, we used populations 2, 4, 9-11, and six other admixed populations (PopA-PopF in [2]) as reference populations using the procedure described in that paper.

### Simulations mimicking European admixture results

Using the approach in [43], we also generated new simulated populations to mimic scenarios that might explain patterns observed in our new analysis of European populations. We used four populations as admixing sources:

a. “Danish”: 200 individuals generated using European cluster C49, where C49 contains 162 Europeans, primarily Danish (Table S3) [10]
b. “German”: the 134 individuals, primarily German, from European cluster C38 [10]
c. “Moroccans”: 25 individuals from Morocco [20, 2]
d. “Evenk”: 12 Evenk from northern Asia [18]

The “Danish” source consisted of a population of 200 individuals generated by intermixing C49 individuals using the approach in [43], with segment sizes (in Morgans) determined by an exponential distribution with rate equal to 200. This was done to mitigate the signal of genuine admixture in C49 (e.g. see Fig 3) from the simulated individuals.

We simulated four different scenarios, each consisting of 50 admixed individuals:

1. “EuroSim1:” 80%/20% of ancestry from Danish/Moroccans, intermixing 100 or 200 generations ago
2. “EuroSim2:” 80%/20% of ancestry from Danish/Evenk, intermixing 100 or 200 generations ago
3. “EuroSim3:” admixture 70 generations ago between Danish/Evenk at 80%/20%, with no subsequent admixture
4. “EuroSim4:” admixture 70 generations ago between Danish/Evenk at 80%/20%, followed by this admixed population intermixing with Germans that contribute 20% of ancestry 10 generations ago (as in Fig S2)

The first of these new European-based simulations is designed to mimic the intermixing of groups related to North Africa (represented by Morocco) and Europe that we observe for several European clusters consisting of Belgians, French and Germans. The remaining three simulations assess our model’s ability to characterise gene flow between Siberian-related groups and Scandinavian populations, which we observe for clusters containing Finns, Norwegians and Swedes, at one or multiple dates. We applied CHROMOPAINTER and fastGLOBETROTTER to each simulation using the same protocol as our real data analysis, though excluding as references for EuroSim1 the Moroccan population used to simulate and excluding as references for EuroSim2-EuroSim4 the Evenk population used to simulate. This reflects a realistic scenario where none of the admixing populations were sampled. For computational simplicity, we used the same paintings of reference populations against each other as was used in the real data analysis, by setting the amount that each other reference population matched to Evenk or Morocco (using either as appropriate) to 0 and re-scaling. This may result in a slight loss in accuracy in these simulations. Results are given in Table 1.

### Analysis of Simulations

For the simulations from [2], we applied GLOBETROTTER and fastGLOBETROTTER to CHROMOPAINTER output generated as described in [2]. When analysing each simulation with GLOBETROTTER and fastGLOBETROTTER, we used the default value of five iterations of inferring dates versus inferring admixture proportions and sources, at each iteration removing reference populations inferred to contribute *<*0.5% of ancestry. In most cases, we constructed admixture probability curves by only considering pairs of DNA segments 1-30cM apart, rounding the distances between pairs to the nearest 0.1cM. An exception is our analyses of the Colombian-Han and French-Brahui simulations with admixture 150 generations ago and ≤20 admixed individuals, where we instead only considered DNA segments 1-5cM apart rounded to the nearest 0.05cM, as recommended by [2] when inferred admixture dates are >55 generations ago when using the default values. In all cases, we used 100 bootstrap re-samples of simulated target population individuals to construct confidence intervals for inferred dates. Following [2], a simulation was considered to have no admixture if the floor of the minimum bootstrap date was 1 or if the maximum bootstrap date was ≥ 400, as the former case is consistent with no admixture and the latter is older than what we can reliably infer with these sample sizes.

### Application to European cohort

We explored admixture in 6,209 European individuals sampled from Belgium, Denmark, Finland, France, Germany, Italy, Norway, Poland, Spain and Sweden genotyped on the Illumina Human 660-Quad SNP array [10]. As reference populations, we used 4,309 individuals sampled from 162 worldwide populations genotyped on a similar platform [10, 16, 2, 17, 18, 19, 20, 21, 22, 23, 24, 25] (Table S4). Different datasets were merged using PLINK [45], after which SNPs with minor-allele-frequency *<*1% or missingness >10% were removed, resulting in 477,417 autosomal SNPs for analysis.

As the precise origins of the 6,209 Europeans were unknown beyond the country level, we clustered them into genetically homogeneous groups prior to inferring admixture. To do so, we first phased individuals jointly using SHAPEIT [46] with the build 36 genetic map. Next, we ran CHROMOPAINTER to form (“paint”) the two phased haploids from each of the 10,522 total individuals as a mosaic of those from all other 10,521 individuals. In particular we estimated the genome-wide average switch (-n flag) and global emission rate (-M flag) by applying 10 iterations of the CHROMOPAINTER expectation-maximization algorithm to paint the phased haploids of 1,052 individuals across chromosomes {4, 10, 15, 22}, painting only one-tenth of individuals and four chromosomes for computational simplicity. We then averaged the estimated values of switch and emission rates across these chromosomes and individuals, giving 52.82727348 and 0.000134461, respectively, and ran CHROMOPAINTER on each of the 10,522 individuals using these fixed values. We applied fineSTRUCTURE [13] to this CHROMOPAINTER output, clustering the 6209 Europeans while fixing the other 162 reference populations as “super-populations” (-f switch) and using 5 million Markov-Chain-Monte-Carlo (MCMC) iterations. FineSTRUCTURE sampled one MCMC iteration out of every 10000, selected the sample from among these with the highest posterior probability, and used 10,000 additional optimisation steps under a greedy approach to find a clustering solution with higher posterior probability [13]. This procedure assigned the 6,209 Europeans into 91 clusters, which fineSTRUCTURE then merged hierarchically, two-at-a-time, under a greedy approach until only two clusters remained. Based on a visual inspection of the tree, we merged a few smaller clusters into large ones, giving 86 clusters that we analysed in subsequent analyses. Cluster names, sample sizes, and population descriptions are provided in Table S3.

To detect admixture events, we applied fastGLOBETROTTER separately to each of these 86 European clusters, using the 162 reference populations as surrogates to the putative admixing sources. To do so, we used CHROMOPAINTER to form the phased haploids of individuals in all 86 clusters and 162 surrogate (reference) populations as mosaics of those from the 162 reference populations. When doing so, we used the same fixed switch and emission parameters in the fineSTRUCTURE analysis for computational convenience. For each of the 86+162 populations, we tabulated the average proportion of genome-wide DNA that individuals from that population match to any haploid in each of the 162 reference populations. Note that an individual is not allowed to match to themselves, so each person in a surrogate population matches to one fewer member from their own population than is the case with individuals in all other surrogate populations and the 86 European clusters. This assymetry may influence inference of the sources involved in the admixture event, though previous analyses have not found this to have a noticeable effect when using similar sample sizes [2, 47]. An exception is surrogate populations with small sample sizes (e.g. ≤2 individuals), which may be upweighted as an ancestry contributer relative to surrogate populations with larger sample sizes [47].

For each haploid from the 86 European clusters, we also used CHROMOPAINTER to generate ten stochastic samples of their mosaic matching to haploids from the 162 reference populations. As in GLOBETROTTER, fastGLOBETROTTER takes as input both the mosaic painting samples of all target population individuals and the average proportion of DNA that the target and 162 surrogate populations matches to each of the 162 reference populations, using these to infer dates and infer admixture sources/proportions [2], respectively, in an iterative manner. We used five iterations of alternating between inferring dates and inferring admixture proportions and sources, while removing reference populations inferred to contribute *<*0.5% of ancestry at each iteration. We also used the “null individual” analysis (i.e. null.ind=1) that adjusts date inference for the effects of a post-admixture bottleneck in the target population. We used 100 bootstrap re-samples of individuals’ chromosomes to construct 95% confidence intervals for the inferred admixture date(s). Dates are inferred in generations (*g*), which were converted to years (*y*) using the formula *y* = 1960 − 28 * (*g* + 1), which assumes a generation time of 28 years [26] and an average birthplace of 1960 for sampled individuals.

## Data Access

All data used in this paper are previously published, from https://ega-archive.org/datasets/EGAD00000000120, https://hagsc.org/hgdp/files.html, http://www.evolutsioon.ut.ee/MAIT/caucasus_data/, https://www.ncbi.nlm.nih.gov/geo/query/acc.cgi?acc=GSE22494, https://www.ncbi.nlm.nih.gov/geo/query/acc.cgi?acc=GSE21478, https://www.sanger.ac.uk/resources/downloads/human/hapmap3.html, https://data.mendeley.com/datasets/ckz9mtgrjj/3.

The software fastGLOBETROTTER is available at https://github.com/sahwa/fastGLOBETROTTER, including a detailed tutorial with example files.

## Acknowledgements

This work was supported by a Sir Henry Dale Fellowship jointly funded by the Wellcome Trust and the Royal Society (Grant Number 098386/Z/12/Z) and supported by the National Institute for Health Research University College London Hospitals Biomedical Research Centre (GH), and by a Royal Thai Government Scholarship (PW).

