## Supplementary Information for "An efficient method to identify, date and describe admixture events using haplotype information"

### Supplementary Tables

| sim groups | gen | true % | n | c | $R_1$ | $R_2$ | $FQ_1$ | date | % | source 1 | source 2 |
|---|---|---|---|---|---|---|---|---|---|---|---|
| Brahui/Yoruba | 7 | 5 | 20 | 1 | 0.93 | 0.03 | 1 | 9 (8-11) | 6 | BantuKenya(0.38)BantuSouthAfrica(0.28)Mandenka(0.25) | Balochi(0.85)Makrani(0.15) |
| Brahui/Yoruba | 7 | 20 | 20 | 1 | 0.96 | 0.01 | 1 | 8 (7-9) | 20 | Mandenka(0.31)BantuKenya(0.3)BantuSouthAfrica(0.25) | Balochi(0.85)Makrani(0.15) |
| Brahui/Yoruba | 7 | 50 | 20 | 1 | 0.97 | 0.02 | 1 | 8 (6-9) | 48 | Mandenka(0.33)BantuKenya(0.3)BantuSouthAfrica(0.24) | Balochi(0.89)Makrani(0.11) |
| Brahui/Yoruba | 30 | 5 | 20 | 1 | 0.94 | 0.03 | 1 | 30 (28-32) | 6 | BantuKenya(0.36)Mandenka(0.27)BantuSouthAfrica(0.22) | Balochi(0.89)Makrani(0.11) |
| Brahui/Yoruba | 30 | 20 | 20 | 1 | 0.95 | 0.04 | 1 | 29 (27-31) | 20 | Mandenka(0.31)BantuKenya(0.29)BantuSouthAfrica(0.26) | Balochi(0.88)Makrani(0.12) |
| Brahui/Yoruba | 30 | 50 | 20 | 1 | 0.94 | 0.01 | 1 | 32 (30-34) | 47 | Mandenka(0.34)BantuKenya(0.3)BantuSouthAfrica(0.24) | Balochi(0.92)Makrani(0.08) |
| Brahui/Yoruba | 150 | 5 | 20 | 1 | 0.24 | 0.04 | 1 | 117 (87-160) | 8 | Mandenka(0.2)BantuKenya(0.18)Makrani(0.14) | Balochi(0.81)Makrani(0.1) |
| Brahui/Yoruba | 150 | 20 | 20 | 1 | 0.29 | 0.04 | 1 | 133 (95-159) | 22 | Mandenka(0.25)BantuKenya(0.22)BantuSouthAfrica(0.2) | Balochi(0.82)Makrani(0.11) |
| Brahui/Yoruba | 150 | 50 | 20 | 1 | 0.3 | 0.05 | 1 | 129 (102-154) | 46 | Mandenka(0.32)BantuKenya(0.29)BantuSouthAfrica(0.23) | Balochi(0.76)Makrani(0.1) |
| Yoruba/French | 7 | 5 | 20 | 1 | 0.98 | 0.02 | 1 | 8 (6-10) | 7 | Lithuanian(0.87)BantuKenya(0.06) | Mandenka(0.32)BantuKenya(0.28)BantuSouthAfrica(0.21) |
| Yoruba/French | 7 | 20 | 20 | 1 | 1 | 0.03 | 1 | 7 (6-9) | 25 | Scottish(0.9)Lithuanian(0.07) | Mandenka(0.33)BantuKenya(0.3)BantuSouthAfrica(0.22) |
| Yoruba/French | 7 | 50 | 20 | 1 | 1 | 0.06 | 1 | 7 (6-8) | 46 | Mandenka(0.34)BantuKenya(0.31)BantuSouthAfrica(0.23) | Welsh(0.42)English(0.17)Scottish(0.11) |
| Yoruba/French | 30 | 5 | 20 | 1 | 0.97 | 0.08 | 1 | 29 (25-32) | 7 | Lithuanian(0.9) | Mandenka(0.31)BantuKenya(0.28)BantuSouthAfrica(0.21) |
| Yoruba/French | 30 | 20 | 20 | 1 | 0.99 | 0.05 | 1 | 30 (28-33) | 26 | Scottish(0.58)Ireland(0.3)Lithuanian(0.08) | Mandenka(0.33)BantuKenya(0.3)BantuSouthAfrica(0.23) |
| Yoruba/French | 30 | 50 | 20 | 1 | 0.99 | 0.13 | 1 | 29 (26-31) | 47 | Mandenka(0.33)BantuKenya(0.3)BantuSouthAfrica(0.23) | Welsh(0.45)Scottish(0.15)NorthItalian(0.11) |
| Yoruba/French | 150 | 5 | 20 | 1 | 0.54 | 0.02 | 0.98 | 128 (127-153) | 12 | Lithuanian(0.62)BantuKenya(0.21)BantuSouthAfrica(0.14) | Mandenka(0.33)BantuKenya(0.25)BantuSouthAfrica(0.2) |
| Yoruba/French | 150 | 20 | 20 | 1 | 0.82 | 0.06 | 1 | 154 (128-175) | 21 | Ireland(0.88)Lithuanian(0.11) | Mandenka(0.31)BantuKenya(0.26)BantuSouthAfrica(0.21) |
| Yoruba/French | 150 | 50 | 20 | 1 | 0.85 | 0.07 | 1 | 144 (131-172) | 46 | Mandenka(0.32)BantuKenya(0.29)BantuSouthAfrica(0.23) | Welsh(0.35)Scottish(0.18)Tuscan(0.11) |
| Colomb/Han | 7 | 5 | 7 | 1 | 0.88 | 0.05 | 1 | 8 (6-10) | 25 | HanNchina(0.63)Saudi(0.07)Maya(0.06) | Maya(0.61)Karitiana(0.07)Pima(0.06) |
| Colomb/Han | 7 | 20 | 7 | 1 | 0.97 | 0.2 | 1 | 6 (5-7) | 40 | Tujia(0.79) | Maya(0.6)Karitiana(0.07)Pima(0.06) |
| Colomb/Han | 7 | 50 | 7 | 1 | 0.98 | 0.09 | 1 | 8 (7-8) | 31 | Maya(0.71)Karitiana(0.09)Pima(0.08) | Tujia(0.47)HanNchina(0.11)Tu(0.08) |
| Colomb/Han | 30 | 5 | 7 | 1 | 0.61 | 0.12 | 0.98 | 18 (16-31) | 24 | Tujia(0.36)Maya(0.14)Scottish(0.12) | Maya(0.56)Karitiana(0.07)Pima(0.05) |
| Colomb/Han | 30 | 20 | 7 | 1 | 0.91 | 0.05 | 1 | 32 (30-34) | 37 | Tujia(0.7)Oroqen(0.06) | Maya(0.59)Karitiana(0.07)Pima(0.06) |
| Colomb/Han | 30 | 50 | 7 | 1 | 0.97 | 0.27 | 1 | 33 (32-34) | 31 | Maya(0.69)Karitiana(0.09)Pima(0.08) | Tujia(0.47)HanNchina(0.16)Xibo(0.07) |
| Colomb/Han† | 150 | 5 | 7 | NA | 0.09 | 0.04 | 0.95 | 49 (3-132) | 47 | Maya(0.53)Pima(0.09) | Uzbekistani(0.38)Maya(0.29)Dai(0.07) |
| Colomb/Han† | 150 | 20 | 7 | 1 | 0.14 | 0.04 | 0.99 | 112 (96-224) | 41 | Tujia(0.55)Myanmar(0.19)Karitiana(0.05) | Maya(0.57)Pima(0.07)Pathan(0.06) |
| Colomb/Han† | 150 | 50 | 7 | 1 | 0.34 | 0.02 | 1 | 107 (82-175) | 37 | Maya(0.56)Karitiana(0.07)Scottish(0.06) | Tujia(0.3)HanNchina(0.21)Xibo(0.09) |
| Brahui/Han | 7 | 5 | 20 | 1 | 0.96 | 0.16 | 0.99 | 10 (6-13) | 13 | Tujia(0.8)Dai(0.07) | Balochi(0.3)Pathan(0.14)Iranian(0.14) |
| Brahui/Han | 7 | 20 | 20 | 1 | 0.98 | 0.34 | 1 | 7 (5-9) | 30 | Tujia(0.53)Dai(0.11)Yi(0.06) | Balochi(0.35)Makrani(0.11)Sindhi(0.11) |
| Brahui/Han | 7 | 50 | 20 | 1 | 0.99 | 0.17 | 1 | 9 (8-11) | 50 | Tujia(0.64)Dai(0.11) | Balochi(0.41)Makrani(0.1)Armenian(0.07) |
| Brahui/Han | 30 | 5 | 20 | 1 | 0.92 | 0.17 | 0.99 | 26 (22-34) | 14 | Tujia(0.66)Dai(0.06)Daur(0.06) | Balochi(0.35)Pathan(0.13)Sindhi(0.09) |
| Brahui/Han | 30 | 20 | 20 | 1 | 0.97 | 0.18 | 1 | 32 (27-37) | 29 | Tujia(0.32)Dai(0.18)Miao(0.1) | Balochi(0.4)Makrani(0.1)Sindhi(0.1) |
| Brahui/Han | 30 | 50 | 20 | 1 | 0.97 | 0.04 | 1 | 33 (30-38) | 50 | Tujia(0.45)Dai(0.12)Yi(0.08) | Balochi(0.46)Makrani(0.09)Cypriot(0.06) |
| Brahui/Han | 150 | 5 | 20 | NA | 0.2 | 0.13 | 0.99 | 22 (4-262) | 47 | Balochi(0.68)Makrani(0.1) | Iranian(0.36)Pathan(0.19)Sindhi(0.1) |
| Brahui/Han | 150 | 20 | 20 | 1 | 0.48 | 0.06 | 1 | 123 (89-160) | 32 | Tujia(0.48)Dai(0.07)HanNchina(0.07) | Balochi(0.46)Sindhi(0.08)Makrani(0.08) |
| Brahui/Han | 150 | 50 | 20 | 1 | 0.46 | 0.03 | 1 | 161 (113-215) | 48 | Tujia(0.68)Dai(0.09) | Balochi(0.38)Makrani(0.09)Sindhi(0.09) |
| French/Brahui | 7 | 5 | 20 | 1 | 0.65 | 0.04 | 0.99 | 11 (7-18) | 7 | Balochi(0.69)Georgian(0.15)Tunisian(0.06) | English(0.31)Scottish(0.24)NorthItalian(0.11) |
| French/Brahui | 7 | 20 | 20 | 1 | 0.91 | 0.04 | 1 | 8 (6-10) | 19 | Balochi(0.99) | English(0.31)Welsh(0.2)Scottish(0.16) |
| French/Brahui | 7 | 50 | 20 | 1 | 0.95 | 0.03 | 1 | 7 (5-8) | 48 | English(0.33)Welsh(0.22)Scottish(0.2) | Balochi(0.87)Makrani(0.13) |
| French/Brahui | 30 | 5 | 20 | 1 | 0.56 | 0.03 | 1 | 27 (19-37) | 7 | Balochi(0.78)Georgian(0.09)UAE(0.07) | English(0.38)Welsh(0.19)Scottish(0.12) |
| French/Brahui | 30 | 20 | 20 | 1 | 0.83 | 0.03 | 1 | 31 (26-37) | 19 | Balochi(0.93)Makrani(0.07) | English(0.31)Welsh(0.2)Scottish(0.14) |
| French/Brahui | 30 | 50 | 20 | 1 | 0.91 | 0.02 | 1 | 28 (25-34) | 48 | English(0.29)Scottish(0.22)Welsh(0.17) | Balochi(0.9)Makrani(0.1) |
| French/Brahui† | 150 | 5 | 20 | NA | 0.02 | 0.02 | 0.62 | 66 (1-716) | 31 | Balochi(0.25)IndianJew(0.19)Palestinian(0.19) | English(0.27)Polish(0.19)Norwegian(0.18) |
| French/Brahui† | 150 | 20 | 20 | NA | 0.08 | 0.02 | 0.94 | 161 (84-415) | 41 | Balochi(0.56)Makrani(0.24)BantuKenya(0.09) | Norwegian(0.3)Lithuanian(0.29)Basque(0.18) |
| French/Brahui† | 150 | 50 | 20 | 1 | 0.1 | 0.01 | 1 | 77 (40-206) | 49 | Scottish(0.27)English(0.15)Welsh(0.12) | Balochi(0.71)Makrani(0.12)NorthItalian(0.07) |

Continued on next page

**Table S1 – continued from previous page**

| sim groups | gen | true % | n | c | $R_1$ | $R_2$ | $FQ_1$ | date | % | source 1 | source 2 |
|---|---|---|---|---|---|---|---|---|---|---|---|
| French/Brahui | 150 | 50 | 50 | 1 | 0.36 | 0.04 | 1 | 108 (48-152) | 48 | English(0.18)Scottish(0.15)WestSicilian(0.12) | Balochi(0.64)NorthItalian(0.14)Welsh(0.1) |
| French/Brahui | 150 | 50 | 100 | 1 | 0.6 | 0.14 | 1 | 146 (112-191) | 44 | Balochi(0.83)Makrani(0.11) | Welsh(0.18)English(0.15)Scottish(0.13) |

Table S1: Admixture inference for simulations in [1] (summarised in Fig 2 of main text). Columns give: **sim groups**: the admixing populations (Colomb=Colombian), **gen**: the true date of admixture (generations ago), **true %**: the true proportion of admixture from the second group, **n**: sample size, **c**: fastGLOBETROTTER's conclusion (1 = one admixture date, NA = no admixture), $R_1$: r-squared fit of a single date (i.e. measuring fit of green line to black lines in Fig S6); $R_2$: additional r-squared explained by adding a second date, $FQ_1$: fit quality of a single date between only two sources, with low values ($< 0.975$) indicating $>$2 admixing sources (see [1] for details), **date**: inferred date and 95% CI, %: proportion of admixture from source 1, **source 1:** top three reference populations used in inferred genetic make-up of the minority contributing source (parentheses give the mixture proportions from these references used to describe this source; these values sum to 1 across all reference populations), **source2:** same for majority contributing source. † refers to cases where the date was inferred considering only pairs of DNA segments 1-5cM apart; otherwise pairs of DNA segments 1-30cM apart were analysed (see Methods).

| | | NULL | | | | | non-NULL | | | | |
|---|---|---|---|---|---|---|---|---|---|---|---|
| **gen** | **method** | **c** | **date** | **%** | **source1** | **source2** | **c** | **date** | **%** | **source1** | **source2** |
| **45** | GT | 1 | 47 (45-52) | 45 | Pop10 | Pop2 | 2 | 30 (29-32) | 48 | Pop10 | Pop2 |
| | fastGT | 1 | 47 (43-50) | 45 | Pop10 | Pop2 | 1 | 31 (28-34) | 48 | Pop10 | Pop2 |
| **20** | GT | 1 | 20 (19-21) | 39 | Pop10 | Pop2 | 1 | 17 (16-18) | 44 | Pop10 | Pop2 |
| | fastGT | 1 | 20 (18-21) | 39 | Pop10 | Pop2 | 1 | 18 (15-19) | 44 | Pop10 | Pop2 |
| **10** | GT | 1 | 10 (10-12) | 40 | Pop10 | Pop2 | 1 | 10 (9-11) | 41 | Pop10 | Pop2 |
| | fastGT | 1 | 10 (9-12) | 40 | Pop10 | Pop2 | 1 | 10 (9-11) | 41 | Pop10 | Pop2 |

Table S2: GLOBETROTTER (GT) and fastGLOBETROTTER (fastGT) inference for simulations from [1] that mimic admixture among African ("Pop2") and East Asian ("Pop8") sources, contributing 60% and 40% of ancestry, respectively, followed by a severe bottleneck in the admixed population. See Fig S1 for the demographic history related the admixing populations. **NULL** gives results when using `null.ind=1`, which adjusts date inference for the effects of a post-admixture bottleneck in the target population. **non-NULL** gives results when using `null.ind=0`, which makes no adjustment. Columns give: **gen:** the simulated date of admixture, **c**: GLOBETROTTER/fastGLOBETROTTER's conclusion (1=one admixture date, 2=multiple admixture dates), **date**: inferred date and 95% CI (in generations ago), **%**: proportion of admixture from source 1, **source1:** inferred single surrogate group that best genetically represents the minority contributing source, **source2:** inferred single surrogate group that best genetically represents the majority contributing source.

| pop | cluster make-up | pop | cluster make-up |
|---|---|---|---|
| C1 | Spa(24) Fra(10) Ger(3) Bel(1) Nor(1) | C2 | Spa(88) Fra(5) Bel(2) Ger(1) |
| C3 | Spa(28) Fra(17) Bel(1) | C4 | Spa(54) Fra(21) Ita(1) Swe(1) |
| C5 | Spa(5) Fra(4) | C6 | Ita(23) Bel(1) |
| C7 | Ita(84) Fra(6) Ger(4) Spa(1) Bel(1) | C8 | Ita(45) Ger(2) Bel(1) |
| C9 | Ita(103) Bel(3) Ger(2) Fra(1) Nor(1) | C10 | Swe(10) Fra(4) Bel(3) Ger(3) Den(2) Pol(1) |
| C11 | Swe(13) Ger(9) Ita(1) Fra(1) Bel(1) Den(1) Nor(1) | C12 | Ger(15) Swe(8) Ita(7) Fra(3) Bel(2) Den(1) Nor(1) |
| C13 | Swe(8) Ger(3) Spa(2) Bel(2) Den(1) Nor(1) Fin(1) | C14 | Ita(14) Fra(1) Bel(1) Den(1) Swe(1) |
| C15 | Ita(30) | C16 | Ita(106) Fra(6) Bel(2) Ger(2) |
| C17 | Ita(26) | C18 | Ita(96) Fra(2) |
| C19 | Fra(22) Ita(18) Bel(1) | C20 | Ita(141) Ger(1) |
| C21 | Ita(46) Fra(1) | C22 | Swe(7) Ger(6) |
| C23 | Fin(58) Swe(6) | C24 | Fin(30) Swe(4) |
| C25 | Fin(57) Swe(1) | C26 | Fin(25) Swe(2) |
| C27 | Fin(45) | C28 | Fin(32) |
| C29 | Fin(83) Swe(6) | C30 | Fin(59) Swe(3) |
| C31 | Fin(56) Swe(7) | C32 | Fin(80) Swe(2) |
| C33 | Fin(30) | C34 | Nor(8) Bel(6) Ger(3) Ita(2) Den(2) Swe(2) |
| C35 | Fra(76) Den(1) | C36 | Swe(11) Ger(2) Bel(1) Pol(1) Nor(1) |
| C37 | Pol(54) Ger(17) Swe(8) Fra(3) Ita(1) Bel(1) | C38 | Ger(128) Den(3) Pol(2) Swe(1) |
| C39 | Ger(136) Swe(5) Fra(3) Bel(1) Nor(1) | C40 | Swe(46) Nor(11) Den(8) Ger(4) Fra(2) |
| C41 | Ger(113) Swe(2) Bel(1) | C42 | Ger(82) Den(7) Swe(4) |
| C43 | Ger(43) | C44 | Ger(52) Den(2) Bel(1) |
| C45 | Ger(92) Bel(2) Den(1) Swe(1) | C46 | Ger(111) Bel(5) Fra(2) |
| C47 | Ger(105) Fra(7) Bel(4) Swe(4) Spa(1) Den(1) | C48 | Ger(151) Fra(2) Bel(2) Swe(2) |
| C49 | Den(153) Nor(7) Swe(2) | C50 | Den(140) Swe(12) Nor(2) Ger(1) |
| C51 | Fra(84) Spa(1) Nor(1) Swe(1) | C52 | Fra(191) Bel(7) Ger(5) Swe(2) Spa(1) |
| C53 | Bel(181) Fra(5) Ger(2) Ita(1) | C54 | Bel(203) Ger(1) |
| C55 | Bel(107) | C56 | Swe(23) Nor(9) |
| C57 | Fin(22) Swe(12) | C58 | Swe(15) Fin(2) Den(1) |
| C59 | Swe(44) | C60 | Swe(77) |
| C61 | Swe(71) Nor(2) | C62 | Swe(28) Nor(1) Fin(1) |
| C63 | Swe(88) Nor(2) | C64 | Swe(140) |
| C65 | Swe(211) Nor(1) | C66 | Swe(70) Den(4) |
| C67 | Swe(154) | C68 | Swe(77) Nor(1) |
| C69 | Nor(25) Swe(2) | C70 | Nor(16) |
| C71 | Nor(21) | C72 | Nor(76) Swe(4) |
| C73 | Nor(28) | C74 | Nor(67) |
| C75 | Nor(37) | C76 | Nor(91) Swe(1) |
| C77 | Nor(78) Swe(6) Den(3) | C78 | Nor(41) Swe(1) |
| C79 | Nor(26) Swe(1) | C80 | Nor(86) Swe(2) |
| C81 | Nor(78) Swe(1) | C82 | Nor(53) |
| C83 | Nor(52) Swe(1) | C84 | Nor(58) |
| C85 | Nor(32) | C86 | Nor(36) |

Table S3: Number of individuals from each country in each Europe cluster. Bel=Belgium, Den=Denmark, Fin=Finland, Fra=France, Ger=Germany, Ita=Italy, Nor=Norway, Pol=Poland, Spa=Spain, Swe=Sweden

| pop | $n$ | region | pop | $n$ | region |
|---|---|---|---|---|---|
| colombian(g) | 7 | Americas | karitiana(g) | 11 | Americas |
| maya(g) | 21 | Americas | pima(g) | 14 | Americas |
| surui(g) | 5 | Americas | bantukenya(g) | 11 | Bantu |
| bantusouthafrica(g) | 8 | Bantu | luhya(c) | 94 | Bantu |
| biakapygmy(g) | 21 | Central Africa | hadza(e) | 3 | Central Africa |
| mbutipygmy(g) | 13 | Central Africa | sandawe(e) | 28 | Central Africa |
| balochi(g) | 24 | Central South Asia | bengali(h) | 1 | Central South Asia |
| bhunjia(h) | 1 | Central South Asia | brahmin(h) | 11 | Central South Asia |
| brahui(g) | 25 | Central South Asia | burusho(g) | 25 | Central South Asia |
| chamar(h) | 10 | Central South Asia | chenchu(b) | 4 | Central South Asia |
| dharkar(h) | 8 | Central South Asia | dhurwa(h) | 1 | Central South Asia |
| dusadh(h) | 7 | Central South Asia | gond(h) | 4 | Central South Asia |
| hakkipikki(b) | 3 | Central South Asia | indian(a) | 1 | Central South Asia |
| indianjew(a) | 8 | Central South Asia | kalash(g) | 23 | Central South Asia |
| kanjar(h) | 5 | Central South Asia | karnataka(a) | 8 | Central South Asia |
| kol(h) | 16 | Central South Asia | kshatriya(h) | 7 | Central South Asia |
| kurmi(h) | 1 | Central South Asia | kurumba(h) | 4 | Central South Asia |
| lambadi(h) | 1 | Central South Asia | makrani(g) | 25 | Central South Asia |
| mawasi(h) | 1 | Central South Asia | meena(h) | 1 | Central South Asia |
| meghawal(h) | 1 | Central South Asia | muslim(h) | 5 | Central South Asia |
| nihali(h) | 2 | Central South Asia | pathan(g) | 22 | Central South Asia |
| piramalaikallar(h) | 8 | Central South Asia | sakd(a) | 4 | Central South Asia |
| sindhi(g) | 24 | Central South Asia | tamilnadu(h) | 2 | Central South Asia |
| tharus(h) | 2 | Central South Asia | upcaste(b) | 5 | Central South Asia |
| velamas(h) | 9 | Central South Asia | hazara(g) | 22 | Central South Asia2 |
| kyrgyz(f) | 16 | Central South Asia2 | uygur(g) | 10 | Central South Asia2 |
| uzbekistani(a) | 15 | Central South Asia2 | maasai(c) | 97 | East Africa |
| kumyk(k) | 14 | East Asia | belorussian(a) | 9 | East Europe |
| bulgarian(k) | 31 | East Europe | chuvash(a) | 17 | East Europe |
| croatian(d) | 19 | East Europe | finnish(d) | 2 | East Europe |
| hungarian(a) | 19 | East Europe | lithuanian(a) | 10 | East Europe |
| mordovian(k) | 15 | East Europe | polish(d) | 17 | East Europe |
| romanian(a) | 16 | East Europe | russian(g) | 25 | East Europe |
| ukrainian(k) | 20 | East Europe | ethiopiana(a) | 7 | Ethiopian |
| ethiopianjew(a) | 11 | Ethiopian | ethiopiano(a) | 7 | Ethiopian |
| ethiopiant(a) | 5 | Ethiopian | moroccan(a,d) | 25 | North Africa |
| mozabite(g) | 29 | North Africa | tunisian(a) | 12 | North Africa |
| buryat(i) | 15 | Northeast Asia | daur(g) | 9 | Northeast Asia |
| hezhen(g) | 8 | Northeast Asia | japanese(g) | 28 | Northeast Asia |
| mongolian(g) | 19 | Northeast Asia | oroqen(g) | 9 | Northeast Asia |
| tu(g) | 10 | Northeast Asia | xibo(g) | 9 | Northeast Asia |
| yakut(g) | 25 | Northeast Asia | basque(g) | 24 | Northwest Europe |
| english(d) | 8 | Northwest Europe | french(g) | 28 | Northwest Europe |
| german(d) | 30 | Northwest Europe | germanyaustria(d) | 4 | Northwest Europe |
| irish(d) | 7 | Northwest Europe | norwegian(d) | 18 | Northwest Europe |
| orcadian(g) | 15 | Northwest Europe | scottish(d) | 6 | Northwest Europe |
| spanish(a,d) | 34 | Northwest Europe | welsh(d) | 4 | Northwest Europe |
| MS:NIreland(j) | 61 | Northwest Europe | MS:UK(j) | 1854 | Northwest Europe |
| melanesian(g) | 10 | Oceania | papuan(g) | 17 | Oceania |
| sankhomani(e) | 30 | San | sannamibia(g) | 5 | San |
| altai(i) | 13 | Siberia | burya(i) | 2 | Siberia |
| chukchi(i) | 5 | Siberia | dolgan(i) | 7 | Siberia |
| evenk(i) | 12 | Siberia | ket(i) | 2 | Siberia |
| koryake(i) | 5 | Siberia | nganassan(i) | 10 | Siberia |
| selkup(i) | 10 | Siberia | tuva(i) | 13 | Siberia |
| yukagir(i) | 4 | Siberia | greek(d) | 16 | South Europe |

Continued on next page

| Table S4 – continued from previous page | | | | | |
|---|---|---|---|---|---|
| **pop** | $n$ | **region** | **pop** | $n$ | **region** |
| northitalian(g) | 12 | South Europe | sardinian(g) | 28 | South Europe |
| siciliane(d) | 10 | South Europe | southitalian(d) | 18 | South Europe |
| tsi(c) | 98 | South Europe | tuscan(g) | 8 | South Europe |
| westsicilian(d) | 10 | South Europe | bedouin(g) | 45 | South Middle East |
| egyptian(a) | 12 | South Middle East | jordanian(a) | 20 | South Middle East |
| lebanese(a) | 5 | South Middle East | palestinian(g) | 46 | South Middle East |
| saudi(a) | 19 | South Middle East | syrian(a) | 16 | South Middle East |
| uae(d) | 14 | South Middle East | yemeni(a) | 9 | South Middle East |
| cambodian(g) | 10 | Southeast Asia | dai(g) | 10 | Southeast Asia |
| han(g) | 34 | Southeast Asia | hannchina(g) | 10 | Southeast Asia |
| lahu(g) | 8 | Southeast Asia | malayan(a) | 1 | Southeast Asia |
| miao(g) | 10 | Southeast Asia | myanmar(a) | 3 | Southeast Asia |
| naga(h) | 4 | Southeast Asia | naxi(g) | 8 | Southeast Asia |
| she(g) | 10 | Southeast Asia | tujia(g) | 10 | Southeast Asia |
| yi(g) | 10 | Southeast Asia | mandenka(g) | 22 | West Africa |
| yoruba(g) | 21 | West Africa | abhkasian(k) | 20 | West Asia |
| adygei(g) | 17 | West Asia | armenian(a) | 35 | West Asia |
| balkar(k) | 19 | West Asia | chechen(k) | 20 | West Asia |
| cypriot(a) | 12 | West Asia | druze(g) | 42 | West Asia |
| georgian(a) | 20 | West Asia | iranian(a) | 20 | West Asia |
| kurd(k) | 6 | West Asia | lezgin(a) | 18 | West Asia |
| nogay(k) | 16 | West Asia | northossetian(k) | 15 | West Asia |
| tajik(k) | 15 | West Asia | turkish(a) | 19 | West Asia |
| turkishe(d) | 23 | West Asia | turkishn(d) | 20 | West Asia |
| turkishs(d) | 20 | West Asia | turkmen(k) | 10 | West Asia |

Table S4: Surrogate groups used as proxies to ancestral sources in each European cluster, along with (approximate) regional assignments designated for this paper ($n$=sample size). The published source of each population is given in parentheses, with a=[2], b=[3], c=[4], d=[1], e=[5], f=[6], g=[7], h=[8], i=[9], j=[10], k=[11].

| pop | n | cluster make-up | c | AF/EA | $R_1$ | $R_2$ | $FQ_1$ | date | % | source 1 | source 2 |
|---|---|---|---|---|---|---|---|---|---|---|---|
| C1 | 39 | Spa(24)Fra(10)Ger(3) | 1 | 9.9/0 | 0.96 | 0.32 | 0.98 | 1092 (924-1288) | 0.08 | moroccan(0.34)yoruba(0.29)tsi(0.18) | welsh(0.32)spanish(0.24)MS:NIreland(0.11) |
| C2 | 96 | Spa(88)Fra(5)Bel(2) | M | 12.1/0 | 0.98 | 0.26 | 0.88 | 868 (756-1008) | 0.18<br>0.15 | lithuanian(0.55)mandenka(0.15)yoruba(0.07)<br>basque(0.67)welsh(0.24) | welsh(0.42)basque(0.16)tsi(0.15)<br>spanish(0.49)french(0.26)scottish(0.06) |
| C3 | 46 | Spa(28)Fra(17)Bel(1) | M | 9.5/0 | 0.91 | 0.24 | 0.9 | 616 (448-812) | 0.34<br>0.17 | basque(0.56)french(0.29)welsh(0.15)<br>moroccan(0.4)basque(0.23)tsi(0.11) | english(0.29)welsh(0.12)tsi(0.11)<br>french(0.58)basque(0.17)welsh(0.12) |
| C4 | 77 | Spa(54)Fra(21)Ita(1) | 2 | 6.4/1 | 0.97 | 0.35 | 0.88 | 1344 (1232-1456)<br>364B (784B-308) | 0.21<br>0.07 | tsi(0.38)basque(0.23)welsh(0.19)<br>tsi(0.34)mozabite(0.14)mandenka(0.12) | english(0.27)spanish(0.26)scottish(0.2)<br>french(0.43)spanish(0.2)welsh(0.15) |
| C5 | 9 | Spa(5)Fra(4) | N | 2.6/0 | 0.24 | 0.03 | 1 | 952 (476-1820) | 0.29 | scottish(0.28)french(0.24)tuscan(0.19) | basque(0.68)spanish(0.17)welsh(0.15) |
| C6 | 24 | Ita(23)Bel(1) | M | 8.5/0 | 0.89 | 0.08 | 0.83 | 1008 (784-1344) | 0.28<br>0.39 | welsh(0.3)southitalian(0.27)tsi(0.12)<br>sardinian(0.78)welsh(0.08) | sardinian(0.61)MS:NIreland(0.12)welsh(0.1)<br>sardinian(0.19)welsh(0.18)tsi(0.17) |
| C7 | 96 | Ita(84)Fra(6)Ger(4) | 1 | 35.5/3.1 | 0.99 | 0.27 | 0.98 | 980 (840-1064) | 0.19 | georgian(0.36)welsh(0.2)mandenka(0.19) | tsi(0.2)welsh(0.17)lezgin(0.16) |
| C8 | 48 | Ita(45)Ger(2)Bel(1) | U | 28.1/0 | 0.88 | 0.26 | 0.89 | 812 (476-1148) | 0.23 | armenian(0.46)tsi(0.19)mandenka(0.1) | welsh(0.26)croatian(0.26)tsi(0.1) |
| C9 | 110 | Ita(103)Bel(3)Ger(2) | M | 31.7/0 | 0.97 | 0.25 | 0.92 | 672 (504-840) | 0.23<br>0.39 | armenian(0.43)tsi(0.15)mandenka(0.11)<br>welsh(0.32)tsi(0.16)spanish(0.09) | welsh(0.23)tsi(0.12)norwegian(0.11)<br>greek(0.16)southitalian(0.13)tsi(0.12) |
| C10 | 23 | Swe(10)Fra(4)Bel(3) | 1 | 17.8/0.3 | 0.77 | 0.31 | 0.98 | 700 (280-1036) | 0.26 | tuscan(0.19)moroccan(0.18)cypriot(0.14) | german(0.53)welsh(0.23)polish(0.09) |
| C11 | 27 | Swe(13)Ger(9)Ita(1) | M | 9/0 | 0.71 | 0.05 | 0.96 | 448 (140-756) | 0.43<br>0.26 | welsh(0.28)tsi(0.23)georgian(0.14)<br>welsh(0.58)MS:NIreland(0.15)tsi(0.12) | belorussian(0.69)lithuanian(0.31)<br>croatian(0.75) |
| C12 | 37 | Ger(15)Swe(8)Ita(7) | M | 1.3/2.2 | 0.62 | 0.08 | 0.97 | 1064 (616-1176) | 0.43<br>0.36 | MS:UK(0.58)norwegian(0.12)welsh(0.1)<br>welsh(0.32)syrian(0.12)norwegian(0.08) | ukrainian(0.39)croatian(0.31)romanian(0.12)<br>polish(0.37)lithuanian(0.33)welsh(0.27) |
| C13 | 18 | Swe(8)Ger(3)Spa(2) | 1 | 10.1/5.3 | 0.78 | 0.08 | 0.98 | 1064 (868-1316) | 0.26 | greek(0.58)daur(0.24)welsh(0.09) | croatian(0.25)norwegian(0.16)MS:UK(0.14) |
| C14 | 18 | Ita(14)Fra(1)Bel(1) | M | 28.2/0 | 0.52 | 0.05 | 0.82 | 532 (196-1344) | 0.32<br>0.42 | sardinian(0.62)tsi(0.22)welsh(0.08)<br>tsi(0.21)sardinian(0.17)armenian(0.1) | welsh(0.22)armenian(0.1)german(0.1)<br>welsh(0.4)sardinian(0.2)german(0.14) |
| C15 | 30 | Ita(30) | M | 12.5/0 | 0.76 | 0.04 | 0.86 | 364 (168-980) | 0.47<br>0.23 | tsi(0.76)maasai(0.05)<br>welsh(0.43)MS:NIreland(0.33) | welsh(0.56)MS:NIreland(0.1)sardinian(0.06)<br>tsi(0.65)welsh(0.17)yemeni(0.08) |
| C16 | 116 | Ita(106)Fra(6)Bel(2) | M | 24.2/0 | 0.97 | 0.18 | 0.91 | 504 (336-672) | 0.23<br>0.45 | armenian(0.47)welsh(0.15)tsi(0.1)<br>welsh(0.42)tsi(0.16)german(0.1) | welsh(0.24)MS:NIreland(0.15)lithuanian(0.11)<br>northitalian(0.23)tsi(0.17)kurd(0.15) |
| C17 | 26 | Ita(26) | 1 | 8.8/0.2 | 0.68 | 0.06 | 0.98 | 560 (28-784) | 0.45 | northitalian(0.47)cypriot(0.1)armenian(0.06) | welsh(0.56)tsi(0.18)MS:NIreland(0.14) |
| C18 | 98 | Ita(96)Fra(2) | 1 | 11.9/0 | 0.89 | 0.06 | 0.98 | 476 (336-672) | 0.32 | tsi(0.17)northitalian(0.14)tunisian(0.12) | welsh(0.28)german(0.18)tsi(0.15) |
| C19 | 41 | Fra(22)Ita(18)Bel(1) | 1 | 12.8/0 | 0.84 | 0.04 | 0.98 | 420 (168-672) | 0.24 | greek(0.2)moroccan(0.15)tsi(0.15) | welsh(0.27)english(0.27)german(0.15) |
| C20 | 142 | Ita(141)Ger(1) | M | 11.7/0 | 0.94 | 0.1 | 0.97 | 420 (252-560) | 0.35<br>0.26 | tsi(0.21)southitalian(0.16)turkishe(0.14)<br>MS:UK(0.53)tsi(0.19)german(0.15) | welsh(0.32)scottish(0.15)tsi(0.15)<br>northitalian(0.29)croatian(0.17)welsh(0.16) |
| C21 | 47 | Ita(46)Fra(1) | 1 | 13.3/0 | 0.81 | 0.05 | 0.99 | 308 (84-588) | 0.26 | tsi(0.21)armenian(0.17)jordanian(0.1) | german(0.52)welsh(0.21)tsi(0.13) |
| C22 | 13 | Swe(7)Ger(6) | N | 71.3/0 | 0.14 | 0.03 | 0.89 | 1848 (1064B-1904) | 0.46 | greek(0.5)lezgin(0.24)lithuanian(0.19) | armenian(0.74)tsi(0.08) |
| C23 | 64 | Fin(58)Swe(6) | 1 | 1.1/7.9 | 0.93 | 0.22 | 0.98 | 280 (56-420) | 0.31 | russian(0.62)oroqen(0.16)nganassan(0.13) | welsh(0.5)norwegian(0.36)russian(0.1) |
| C24 | 34 | Fin(30)Swe(4) | 1 | 0/7.6 | 0.89 | 0.24 | 0.99 | 336 (0-560) | 0.34 | russian(0.62)oroqen(0.22)nganassan(0.05) | welsh(0.77)russian(0.11)sardinian(0.1) |
| C25 | 58 | Fin(57)Swe(1) | 1 | 0/8.9 | 0.94 | 0.24 | 0.98 | 28B (168B-140) | 0.34 | russian(0.56)oroqen(0.1)nganassan(0.1) | welsh(0.55)norwegian(0.27)orcadian(0.1) |
| C26 | 27 | Fin(25)Swe(2) | 1 | 0/11.7 | 0.88 | 0.11 | 0.99 | 308 (112-560) | 0.27 | russian(0.58)daur(0.26)oroqen(0.17) | welsh(0.51)norwegian(0.39) |
| C27 | 45 | Fin(45) | 1 | 0/9.7 | 0.92 | 0.13 | 0.99 | 84 (112B-308) | 0.3 | russian(0.51)oroqen(0.19)dolgan(0.18) | welsh(0.53)norwegian(0.39)russian(0.06) |
| C28 | 32 | Fin(32) | 1 | 1/9.6 | 0.88 | 0.24 | 0.99 | 224 (28B-616) | 0.31 | russian(0.51)nganassan(0.1)oroqen(0.09) | welsh(0.62)norwegian(0.26)russian(0.1) |
| C29 | 89 | Fin(83)Swe(6) | M | 0/7.9 | 0.92 | 0.27 | 0.97 | 756 (448-952) | 0.32<br>0.23 | russian(0.6)oroqen(0.21)nganassan(0.07)<br>welsh(0.73)norwegian(0.1)oroqen(0.1) | welsh(0.4)norwegian(0.33)russian(0.17)<br>finnish(0.56)russian(0.37) |
| C30 | 62 | Fin(59)Swe(3) | 1 | 2.4/11 | 0.93 | 0.21 | 0.99 | 56B (224B-224) | 0.29 | russian(0.48)oroqen(0.29)dolgan(0.1) | welsh(0.78)russian(0.15) |
| C31 | 63 | Fin(56)Swe(7) | 1 | 1.8/13.5 | 0.94 | 0.24 | 0.99 | 28 (140B-252) | 0.32 | russian(0.46)oroqen(0.43) | welsh(0.77)russian(0.17) |
| C32 | 82 | Fin(80)Swe(2) | 1 | 0/9.8 | 0.93 | 0.31 | 0.99 | 0 (168B-224) | 0.41 | russian(0.5)yukagir(0.2)koryake(0.12) | welsh(0.75)russian(0.21) |
| C33 | 30 | Fin(30) | 1 | 0/13.3 | 0.88 | 0.13 | 0.99 | 140 (140B-364) | 0.3 | russian(0.43)oroqen(0.37)evenk(0.07) | norwegian(0.41)welsh(0.33)orcadian(0.13) |
| C34 | 23 | Nor(8)Bel(6)Ger(3) | 1 | 2.7/0.3 | 0.71 | 0.09 | 0.99 | 1260 (868-1764) | 0.04 | sardinian(0.48)orcadian(0.34)yoruba(0.07) | welsh(0.6)MS:NIreland(0.2)MS:UK(0.13) |

Continued on next page

Table S5 – continued from previous page

| pop | n | cluster make-up | c | AF/EA | $R_1$ | $R_2$ | $FQ_1$ | date | % | source 1 | source 2 |
|---|---|---|---|---|---|---|---|---|---|---|---|
| C35 | 77 | Fra(76)Den(1) | 1 | 11.5/0.4 | 0.82 | 0.06 | 0.99 | 280 (56B-784) | 0.09 | tsi(0.22)cypriot(0.21)moroccan(0.19) | welsh(0.42)scottish(0.14)irish(0.14) |
| C36 | 16 | Swe(11)Ger(2)Bel(1) | M | 2.6/9.2 | 0.87 | 0.13 | 0.93 | 812 (532-1120) | 0.16<br>0.28 | lithuanian(0.56)miao(0.16)she(0.16)<br>welsh(0.82) | lithuanian(0.27)polish(0.25)welsh(0.21)<br>lithuanian(0.64)russian(0.12)koryake(0.06) |
| C37 | 84 | Pol(54)Ger(17)Swe(8) | M | 1.2/4.6 | 0.77 | 0.04 | 0.86 | 784 (560-924) | 0.25<br>0.15 | welsh(0.91)<br>welsh(0.56)russian(0.16)buryat(0.13) | lithuanian(0.59)belorussian(0.19)mordovian(0.11)<br>polish(0.74)lithuanian(0.18) |
| C38 | 134 | Ger(128)Den(3)Pol(2) | 1 | 0/0 | 0.9 | 0.25 | 1 | 1232 (1148-1344) | 0.43 | welsh(0.42)MS:UK(0.42)tsi(0.1) | belorussian(0.79)polish(0.2) |
| C39 | 146 | Ger(136)Swe(5)Fra(3) | 1 | 0/0 | 0.9 | 0.12 | 1 | 1148 (1036-1260) | 0.49 | belorussian(0.69)polish(0.14)ukrainian(0.11) | MS:UK(0.36)english(0.22)welsh(0.2) |
| C40 | 71 | Swe(46)Nor(11)Den(8) | 1 | 0.4/0.7 | 0.79 | 0.1 | 0.99 | 1064 (896-1232) | 0.3 | belorussian(0.34)ukrainian(0.14)norwegian(0.13) | MS:UK(0.5)welsh(0.29)german(0.11) |
| C41 | 116 | Ger(113)Swe(2)Bel(1) | 1 | 0/0 | 0.86 | 0.17 | 1 | 1316 (1204-1428) | 0.38 | polish(0.64)ukrainian(0.25)lithuanian(0.07) | MS:UK(0.49)welsh(0.26)german(0.2) |
| C42 | 93 | Ger(82)Den(7)Swe(4) | 1 | 0/0 | 0.87 | 0.11 | 1 | 1260 (1176-1372) | 0.37 | polish(0.74)lithuanian(0.07)german(0.06) | MS:UK(0.54)welsh(0.4)german(0.06) |
| C43 | 43 | Ger(43) | 1 | 9.8/0 | 0.74 | 0.04 | 1 | 532 (420-812) | 0.19 | greek(0.2)tsi(0.14)cypriot(0.12) | welsh(0.54)MS:UK(0.21)german(0.13) |
| C44 | 55 | Ger(52)Den(2)Bel(1) | 1 | 4.6/0 | 0.74 | 0.05 | 1 | 588 (364-812) | 0.13 | croatian(0.29)belorussian(0.23)tsi(0.23) | welsh(0.9)norwegian(0.07) |
| C45 | 96 | Ger(92)Bel(2)Den(1) | 1 | 6.2/0 | 0.88 | 0.16 | 1 | 784 (532-896) | 0.18 | greek(0.31)ukrainian(0.13)cypriot(0.12) | welsh(0.64)MS:UK(0.22)norwegian(0.1) |
| C46 | 118 | Ger(111)Bel(5)Fra(2) | 1 | 8.3/0 | 0.91 | 0.08 | 1 | 532 (392-672) | 0.2 | greek(0.22)tsi(0.12)cypriot(0.12) | MS:UK(0.56)welsh(0.27)german(0.1) |
| C47 | 122 | Ger(105)Fra(7)Bel(4) | 1 | 10.5/0 | 0.92 | 0.11 | 1 | 616 (420-700) | 0.25 | cypriot(0.18)greek(0.16)tsi(0.15) | welsh(0.34)MS:UK(0.34)german(0.19) |
| C48 | 157 | Ger(151)Fra(2)Bel(2) | 1 | 5.6/0 | 0.89 | 0.14 | 0.99 | 756 (616-924) | 0.31 | croatian(0.23)greek(0.22)armenian(0.11) | welsh(0.37)MS:UK(0.27)german(0.23) |
| C49 | 162 | Den(153)Nor(7)Swe(2) | 1 | 2.9/0 | 0.89 | 0.09 | 1 | 644 (504-812) | 0.15 | lithuanian(0.29)tsi(0.23)croatian(0.2) | welsh(0.81)MS:NIreland(0.11)norwegian(0.08) |
| C50 | 155 | Den(140)Swe(12)Nor(2) | 1 | 0.9/0.3 | 0.81 | 0.11 | 1 | 896 (812-1092) | 0.2 | polish(0.5)tsi(0.12)ukrainian(0.11) | welsh(0.82)norwegian(0.12)MS:NIreland(0.06) |
| C51 | 87 | Fra(84)Spa(1)Nor(1) | U | 11.8/0 | 0.93 | 0.23 | 0.82 | 392 (224-504) | 0.14 | tsi(0.25)moroccan(0.17)greek(0.09) | french(0.33)welsh(0.26)scottish(0.18) |
| C52 | 206 | Fra(191)Bel(7)Ger(5) | M | 10.8/0 | 0.97 | 0.17 | 0.97 | 420 (308-560) | 0.15<br>0.44 | tsi(0.25)greek(0.21)moroccan(0.16)<br>welsh(0.75)MS:NIreland(0.14) | english(0.38)welsh(0.24)MS:UK(0.11)<br>english(0.34)welsh(0.26)greek(0.12) |
| C53 | 189 | Bel(181)Fra(5)Ger(2) | 1 | 11.6/0 | 0.96 | 0.17 | 1 | 532 (420-616) | 0.16 | tsi(0.19)greek(0.16)cypriot(0.14) | MS:UK(0.67)welsh(0.22)german(0.06) |
| C54 | 204 | Bel(203)Ger(1) | 1 | 8.5/0 | 0.95 | 0.1 | 1 | 504 (392-560) | 0.16 | greek(0.22)armenian(0.22)tsi(0.21) | MS:UK(0.51)welsh(0.34)norwegian(0.06) |
| C55 | 107 | Bel(107) | 1 | 7.9/0 | 0.92 | 0.1 | 1 | 560 (364-672) | 0.15 | greek(0.25)armenian(0.19)tsi(0.18) | MS:UK(0.59)welsh(0.28)norwegian(0.07) |
| C56 | 32 | Swe(23)Nor(9) | 1 | 0/5.6 | 0.89 | 0.18 | 0.99 | 784 (616-952) | 0.25 | russian(0.72)nganassan(0.1)oroqen(0.1) | welsh(0.62)norwegian(0.2)lithuanian(0.11) |
| C57 | 34 | Fin(22)Swe(12) | 1 | 0/4.9 | 0.88 | 0.23 | 0.99 | 476 (224-672) | 0.32 | russian(0.77)dolgan(0.09)oroqen(0.07) | welsh(0.57)norwegian(0.17)lithuanian(0.13) |
| C58 | 18 | Swe(15)Fin(2)Den(1) | 1 | 0/5.4 | 0.7 | 0.14 | 0.98 | 280 (448B-644) | 0.26 | russian(0.68)oroqen(0.08)nganassan(0.06) | welsh(0.56)norwegian(0.27)MS:NIreland(0.06) |
| C59 | 44 | Swe(44) | 1 | 0/3.8 | 0.88 | 0.26 | 1 | 980 (840-1176) | 0.24 | russian(0.41)norwegian(0.32)chuvash(0.07) | welsh(0.57)MS:NIreland(0.25)norwegian(0.11) |
| C60 | 77 | Swe(77) | 1 | 0/3.5 | 0.91 | 0.21 | 0.99 | 896 (700-1064) | 0.19 | russian(0.39)norwegian(0.2)lithuanian(0.17) | welsh(0.56)MS:NIreland(0.17)norwegian(0.12) |
| C61 | 73 | Swe(71)Nor(2) | 1 | 0/5.6 | 0.91 | 0.27 | 1 | 588 (420-812) | 0.19 | russian(0.41)norwegian(0.27)koryake(0.11) | welsh(0.62)MS:NIreland(0.17)norwegian(0.12) |
| C62 | 30 | Swe(28)Nor(1)Fin(1) | 1 | 0/2.4 | 0.8 | 0.19 | 0.99 | 840 (588-1092) | 0.37 | finnish(0.65)russian(0.2) | welsh(0.62)MS:NIreland(0.19)norwegian(0.1) |
| C63 | 90 | Swe(88)Nor(2) | 1 | 0/3.7 | 0.89 | 0.14 | 0.99 | 672 (560-868) | 0.18 | russian(0.31)norwegian(0.3)lithuanian(0.18) | welsh(0.65)norwegian(0.15)MS:NIreland(0.14) |
| C64 | 140 | Swe(140) | 2 | 0/2.9 | 0.93 | 0.37 | 0.99 | 1288 (1148-1568)<br>1400B (1876-308B) | 0.14<br>0.16 | russian(0.51)lithuanian(0.24)norwegian(0.15)<br>lithuanian(0.28)russian(0.21)norwegian(0.19) | welsh(0.52)norwegian(0.21)lithuanian(0.06)<br>welsh(0.66)MS:NIreland(0.18)russian(0.06) |
| C65 | 212 | Swe(211)Nor(1) | 2 | 0/0.5 | 0.94 | 0.46 | 0.99 | 1288 (1120-1624)<br>700B (1036B-28) | 0.32<br>0.05 | polish(0.26)welsh(0.2)norwegian(0.18)<br>koryake(0.48)russian(0.16)nganassan(0.09) | welsh(0.43)MS:UK(0.27)norwegian(0.17)<br>welsh(0.43)MS:UK(0.18)norwegian(0.16) |
| C66 | 74 | Swe(70)Den(4) | 1 | 0.4/0.3 | 0.68 | 0.09 | 1 | 784 (644-1176) | 0.18 | belorussian(0.34)russian(0.19)ukrainian(0.18) | welsh(0.63)norwegian(0.16)MS:UK(0.1) |
| C67 | 154 | Swe(154) | 1 | 0/2.2 | 0.88 | 0.17 | 1 | 532 (364-812) | 0.18 | ukrainian(0.71)russian(0.11)norwegian(0.07) | welsh(0.71)MS:NIreland(0.13)norwegian(0.11) |
| C68 | 78 | Swe(77)Nor(1) | 1 | 0/1.6 | 0.69 | 0.08 | 0.99 | 672 (280-896) | 0.22 | norwegian(0.43)lithuanian(0.28)russian(0.07) | welsh(0.7)norwegian(0.16)MS:NIreland(0.15) |
| C69 | 27 | Nor(25)Swe(2) | 1 | 0.3/3.8 | 0.83 | 0.31 | 1 | 1204 (1120-1456) | 0.16 | russian(0.57)chuvash(0.14)nganassan(0.07) | welsh(0.43)norwegian(0.39)MS:NIreland(0.09) |
| C70 | 16 | Nor(16) | 1 | 0/5.3 | 0.73 | 0.11 | 0.99 | 756 (280-1064) | 0.22 | norwegian(0.37)russian(0.34)oroqen(0.06) | welsh(0.66)MS:NIreland(0.22) |
| C71 | 21 | Nor(21) | 1 | 0/5 | 0.79 | 0.12 | 1 | 952 (504-1092) | 0.19 | norwegian(0.56)russian(0.12)nganassan(0.1) | welsh(0.61)MS:NIreland(0.23)norwegian(0.12) |
| C72 | 80 | Nor(76)Swe(4) | 1 | 0.2/6.3 | 0.92 | 0.26 | 1 | 728 (644-952) | 0.12 | norwegian(0.37)russian(0.27)nganassan(0.12) | welsh(0.57)norwegian(0.19)MS:NIreland(0.18) |
| C73 | 28 | Nor(28) | 1 | 0/4.9 | 0.87 | 0.17 | 1 | 868 (588-1092) | 0.17 | norwegian(0.49)russian(0.22)nganassan(0.1) | welsh(0.55)MS:NIreland(0.28)norwegian(0.14) |
| C74 | 67 | Nor(67) | 1 | 0/3.8 | 0.91 | 0.29 | 1 | 1092 (952-1260) | 0.2 | norwegian(0.52)russian(0.2)nganassan(0.1) | welsh(0.6)MS:NIreland(0.23)norwegian(0.12) |

Continued on next page

**Table S5 – continued from previous page**

| pop | n | cluster make-up | c | AF/EA | $R_1$ | $R_2$ | $FQ_1$ | date | % | source 1 | source 2 |
|---|---|---|---|---|---|---|---|---|---|---|---|
| C75 | 37 | Nor(37) | 1 | 0.3/2.9 | 0.83 | 0.05 | 0.99 | 308 (28-672) | 0.09 | lithuanian(0.35)nganassan(0.17)norwegian(0.1) | welsh(0.61)norwegian(0.2)MS:NIreland(0.14) |
| C76 | 92 | Nor(91)Swe(1) | 1 | 0/3.2 | 0.88 | 0.26 | 0.99 | 672 (448-868) | 0.2 | norwegian(0.61)russian(0.21)oroqen(0.06) | welsh(0.67)MS:NIreland(0.16)norwegian(0.14) |
| C77 | 87 | Nor(78)Swe(6)Den(3) | 1 | 0/3.5 | 0.85 | 0.08 | 0.98 | 560 (336-784) | 0.18 | norwegian(0.59)russian(0.15)dolgan(0.07) | welsh(0.65)MS:NIreland(0.19)norwegian(0.12) |
| C78 | 42 | Nor(41)Swe(1) | 1 | 0/3 | 0.66 | 0.02 | 0.99 | 532 (280-1008) | 0.14 | norwegian(0.49)russian(0.18)selkup(0.14) | welsh(0.53)norwegian(0.44) |
| C79 | 27 | Nor(26)Swe(1) | N | 3.9/0 | 0.42 | 0.04 | 0.9 | 700 (252-952) | 0.44 | welsh(0.91) | welsh(0.43)norwegian(0.33)lithuanian(0.16) |
| C80 | 88 | Nor(86)Swe(2) | M | 0/5.6 | 0.89 | 0.29 | 0.95 | 448 (140-672) | 0.36<br>0.45 | norwegian(0.84)nganassan(0.06)<br>lithuanian(0.9)norwegian(0.1) | welsh(0.94)orcadian(0.06)<br>welsh(0.82)surui(0.12) |
| C81 | 79 | Nor(78)Swe(1) | 1 | 0/3.4 | 0.87 | 0.09 | 0.99 | 532 (308-728) | 0.18 | norwegian(0.66)russian(0.14)nganassan(0.08) | welsh(0.54)MS:NIreland(0.34)norwegian(0.1) |
| C82 | 53 | Nor(53) | M | 0.7/2.5 | 0.62 | 0.13 | 0.9 | 504 (196B-840) | 0.24<br>0.3 | norwegian(0.84)selkup(0.06)<br>welsh(0.42)lithuanian(0.41)norwegian(0.09) | MS:NIreland(0.52)welsh(0.45)<br>welsh(0.95) |
| C83 | 53 | Nor(52)Swe(1) | 1 | 0/4.9 | 0.82 | 0.19 | 0.98 | 252 (0-812) | 0.17 | norwegian(0.79)nganassan(0.19) | welsh(0.64)MS:NIreland(0.2)norwegian(0.15) |
| C84 | 58 | Nor(58) | U | 1.3/1.4 | 0.73 | 0.05 | 0.93 | 252 (84-644) | 0.04 | biakapygmy(0.23)yakut(0.18)kalash(0.18) | welsh(0.62)norwegian(0.23)MS:NIreland(0.09) |
| C85 | 32 | Nor(32) | 1 | 0/2.8 | 0.63 | 0.02 | 0.99 | 392 (56B-952) | 0.17 | norwegian(0.84)yakut(0.09) | welsh(0.84)orcadian(0.06)norwegian(0.05) |
| C86 | 36 | Nor(36) | M | 0/3.1 | 0.68 | 0.06 | 0.97 | 392 (28-756) | 0.31<br>0.42 | norwegian(0.81)russian(0.06)oroqen(0.06)<br>welsh(0.79)surui(0.09)russian(0.06) | welsh(0.95)orcadian(0.05)<br>norwegian(1) |

Table S5: Admixture inference for each European cluster. Columns give: **n**: sample size, **cluster make-up**: the three countries most represented in the cluster (counts in parentheses; see Table S3 for full list), **c**: fastGLOBETROTTER's conclusion (1=one admixture date, 2=multiple admixture dates, M=one-date with $>$2 sources, U=admixture difficult to classify, N=caution should be used due to low signal with $R_1 < 0.5$), **AF/EA**: total proportion of DNA matched to *"African/W.Asia"* and *"E.Asia/Siberia"* groups described in Fig 3; $R_1$: r-squared fit of a single date (i.e. measuring fit of green line to black lines in Fig S6); $R_2$: additional r-squared explained by adding a second date, $FQ_1$: fit quality of a single date between only two sources, with low values ($< 0.975$) indicating $>$2 admixing sources (see [1] for details), **date**: inferred date and 95% CI (B=BCE, otherwise CE), %: proportion of admixture from source 1, **source 1:** top three reference populations used in inferred genetic make-up of the minority contributing source (parentheses give the mixture proportions from these references used to describe this source; these values sum to 1 across all reference populations), **source2:** same for majority contributing source.

### Supplementary Figures

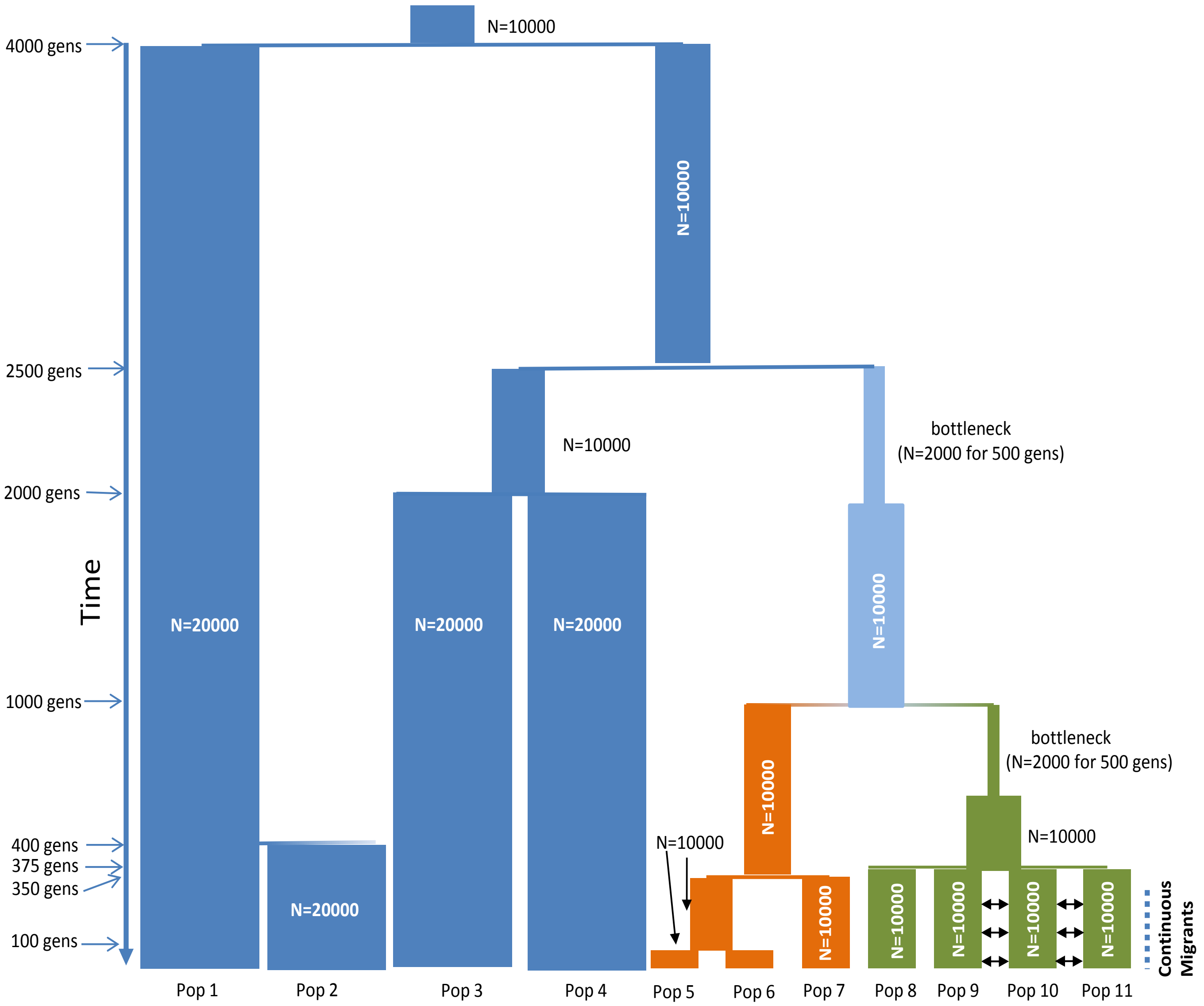


Figure S1: Simulated history for populations 1-11, generated using the coalescent-based software MaCS [12], taken from [1]. Roughly speaking, populations 1-4 in blue are meant to represent diversity in African groups, with populations 5-7 in orange and 8-11 in green representing Western Eurasian and East Asian groups, respectively. 100 generations (gens) denotes the split between Pop5 and Pop6; 350 gens the split between Pop7 and Pop5/Pop6; 375 gens the simultaneous split of populations 8-11; 400 gens the split of Pop1 and Pop2; 1000 gens the split of Pop5/Pop6/Pop7 and Pop8/Pop9/Pop10/Pop11; 2000 gens the split of Pop3 and Pop4; 2500 gens the split of Pop3/Pop4 and Pop5/Pop6/Pop7/Pop8/Pop9/Pop10/Pop11; and 4000 gens the split of Pop1/Pop2 and all other populations.

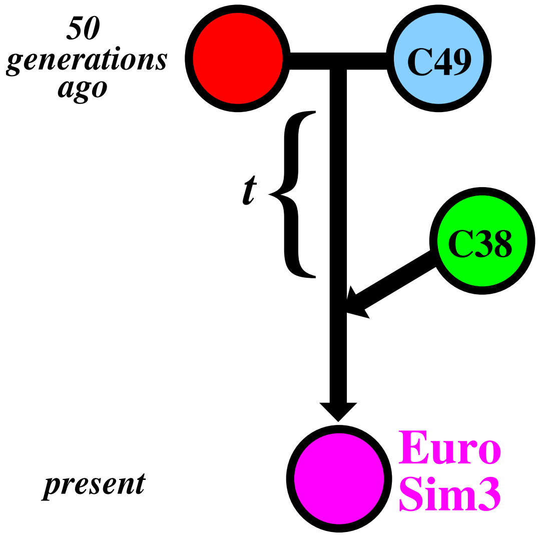


Figure S2: Simulated history for "EuroSim3" (represented by the pink circle) designed to mimic a scenario that might explain patterns observed in our European analysis. Here we simulate admixture occurring 70 generations ago between the Evenk from Northern Asia (red circle) and Europe cluster 49 primarily comprised of Danish (cyan circle), with this admixed group subsequently intermixing with Europe cluster 38 primarily comprised of Germans (green circle) $t$ generations later. Given the genetic similarity between Europe clusters 38 and 49, for small $t$ (i.e. $t < 50$) fastGLOBETROTTER may infer only single pulse of admixture in "EuroSim3", with inferred date somewhere along the branch indicated by $t$ and a higher proportion of European-like ancestry relative to a scenario where there is no additional pulse of admixture from Europe cluster 38.

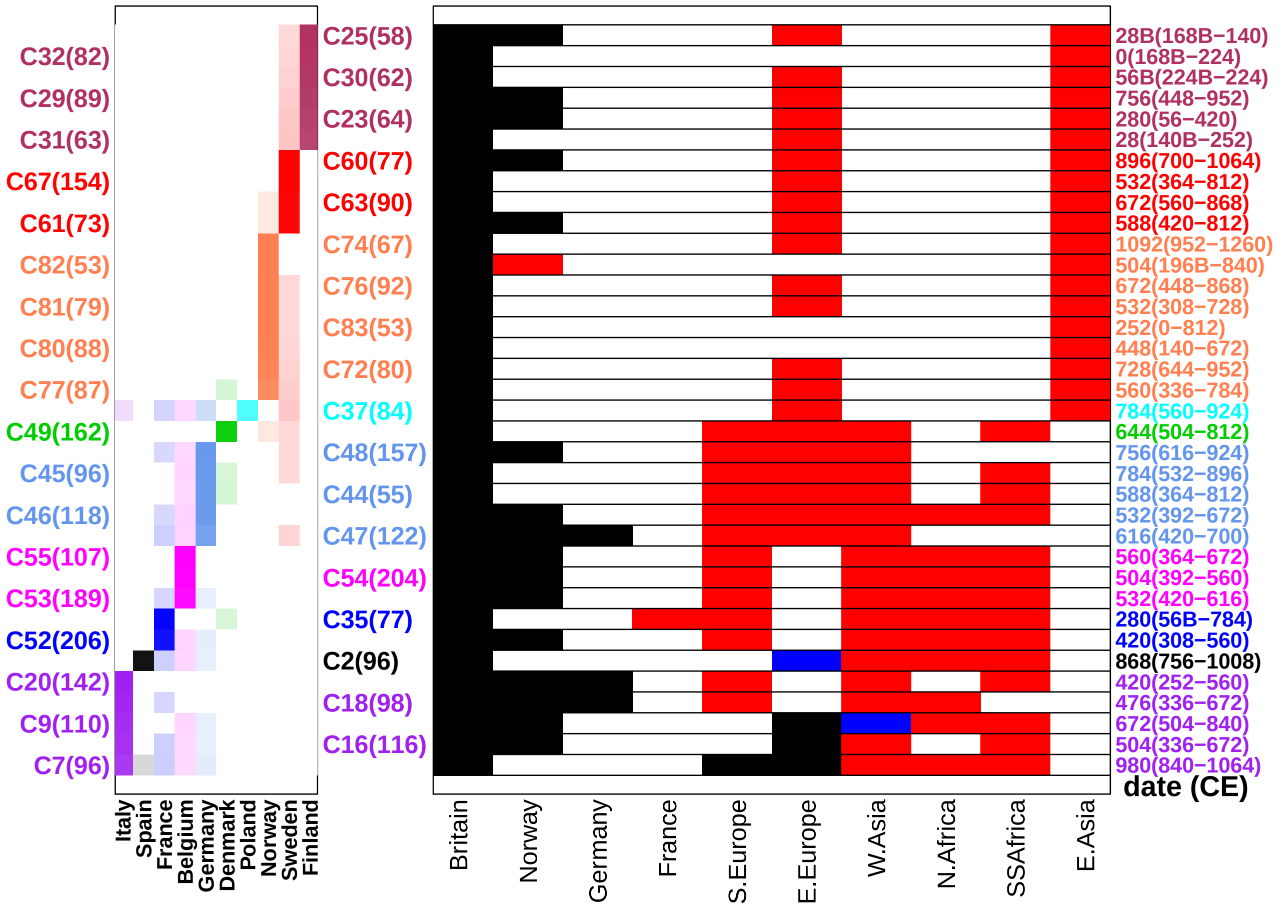


Figure S3: **(right:)** Inferred admixing sources for the 36 Europe clusters that contain $>$50 individuals, have a single inferred date, and have a $>$2% inferred contribution from reference groups in E.Asia/Siberia or in Africa/W.Asia (as in main text Figure 3). Black, red and blue colors refer to distinct admixing sources in each row. Geographic regions (columns) were classified into sources by finding the admixture probability curve with the highest r-squared fit assuming one date of admixture (i.e. fit of green line to black lines in Figures S6-S16), among all curves containing a population from Britain and a population from the given column. If this curve was monotonically decreasing, the region was assigned to the black source, otherwise it was assigned to the red source. In clusters where fastGLOBETROTTER inferred $>$2 sources mixing at the same time, the blue source reflect cases where the fitted line assuming one date (green lines in Figures S6-S16) and the fitted line assuming one date with only two sources (cyan lines in Figures S6-S16) are not both increasing or both decreasing. Britain={MS:UK, english, NIreland, orcadian, scottish, welsh}, Germany={germanyaustria, german}, W.Asia={South Middle East, West Asia}, SSAfrica={West Africa, San, Central Africa, East Africa, Bantu, Ethiopian}, E.Asia={Siberia, Northeast Asia, Southeast Asia}. **(left:)** The proportion of individuals in each cluster (row) from each country (column), with clusters' sample sizes in parentheses and countries' colors based on the map in Figure 3 of the main text. Each cluster's label at left and inferred admixture date (+95% CI) at right is colored according to its majority contributing country.

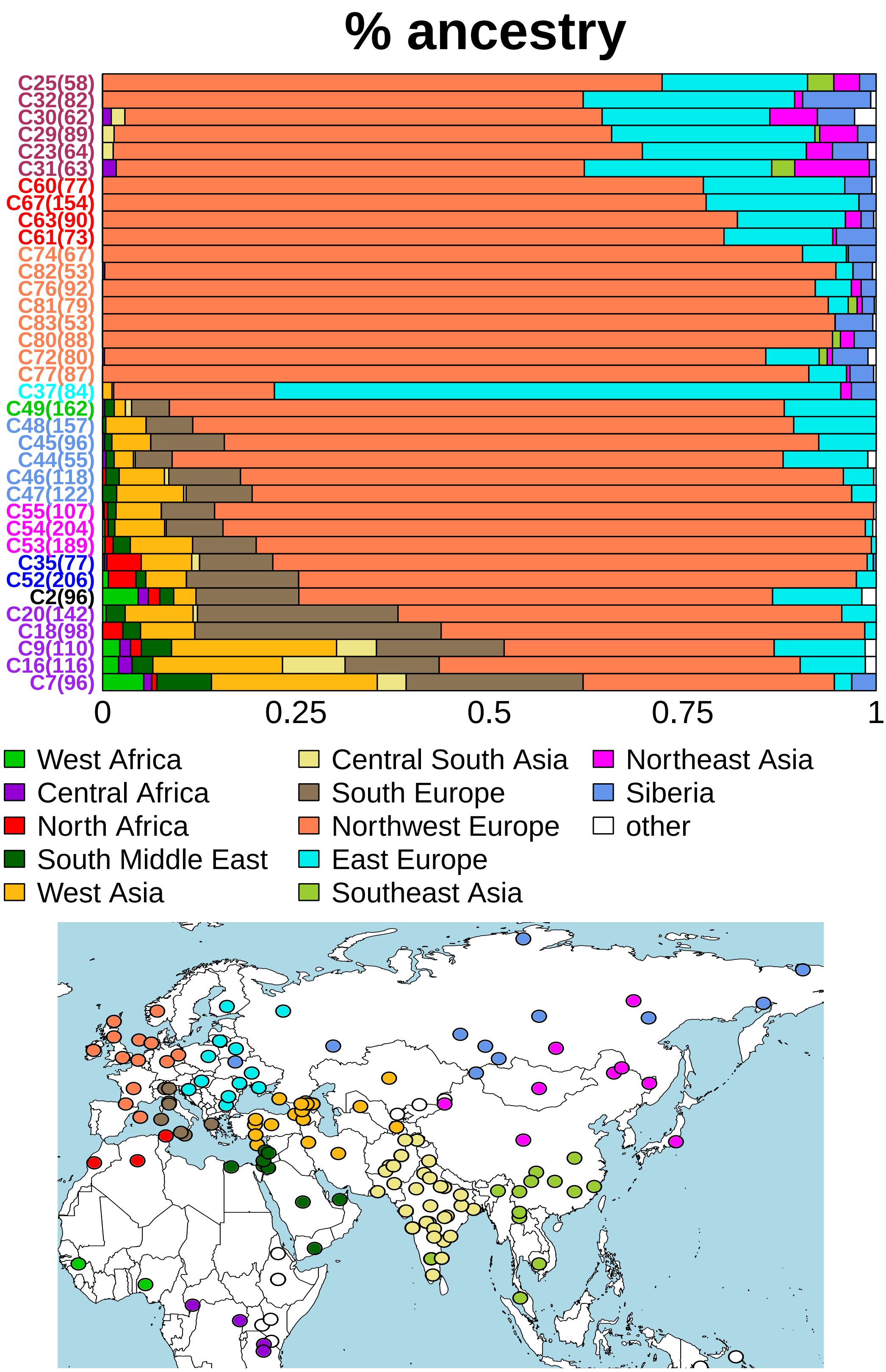


Figure S4: Inferred average proportion of matching to reference populations (bottom map), for all 36 Europe clusters (rows) that contain >50 individuals, have a single inferred date, and have a >2% inferred contribution from reference groups in E.Asia/Siberia or in Africa/W.Asia (as in main text Figure 3).

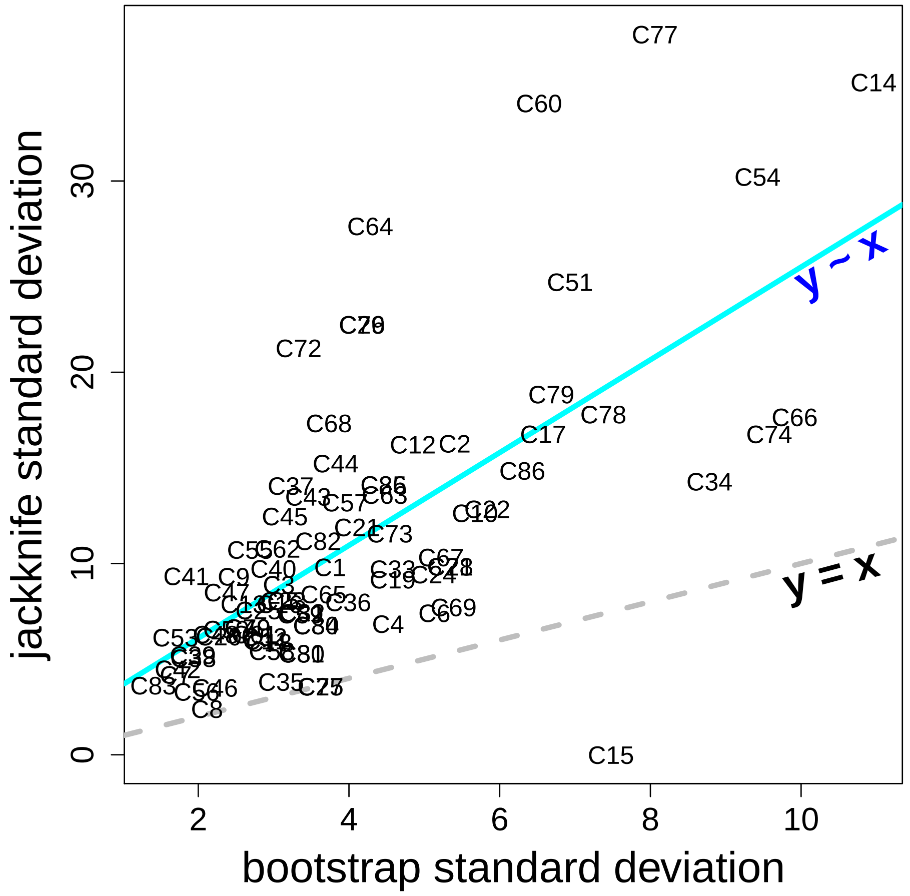


Figure S5: Comparison of standard deviations for date estimates when using jack-knifing versus bootstraping, for all 77 European clusters that inferred “one-date” or “one-date, multiway” admixture. The cyan line depicts the regression line, while the dotted grey line depicts the line of equality. Jack-knife estimates are consistently larger than bootstrap estimates, as expected [13], but they are highly correlated.

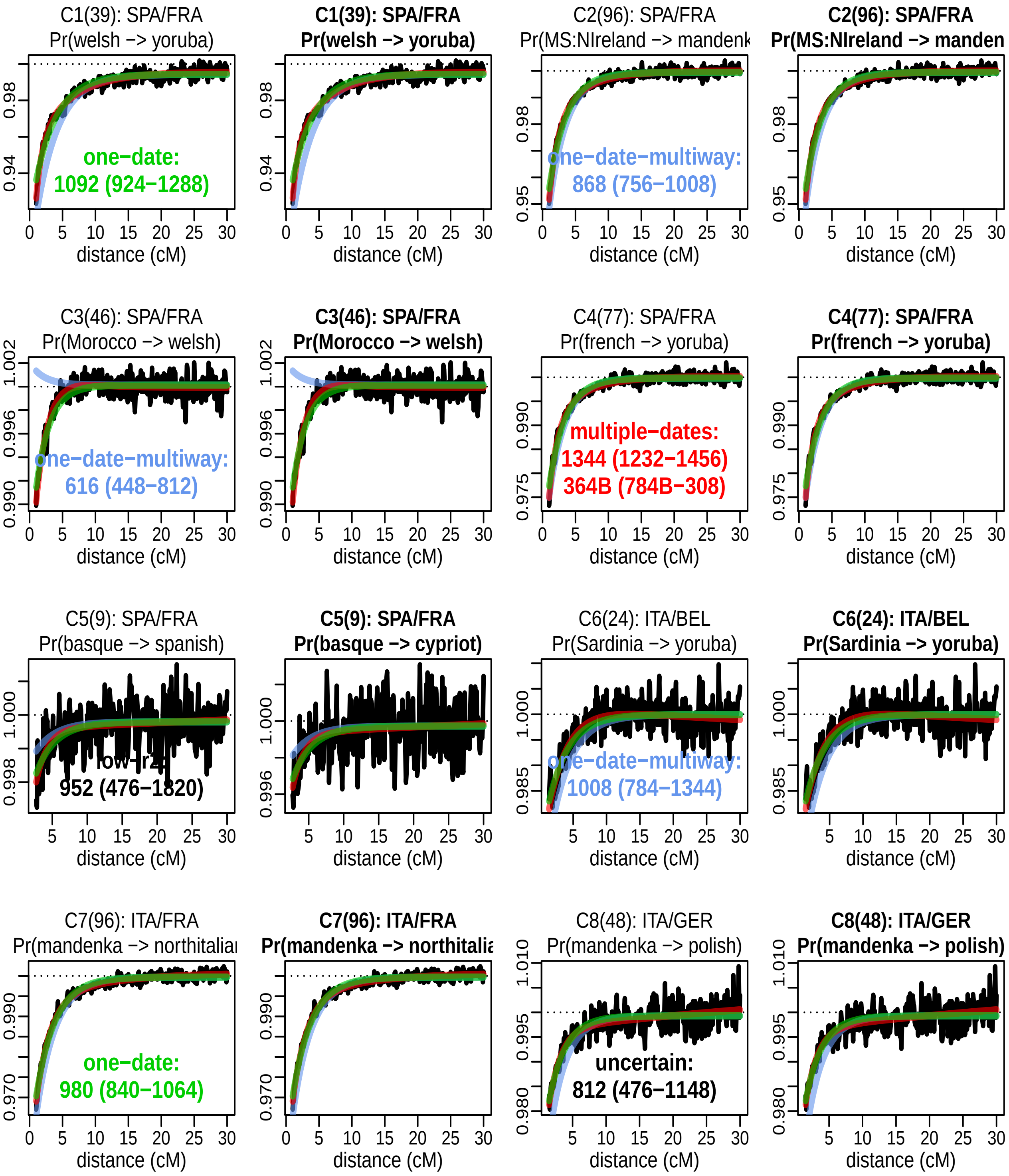


Figure S6: FastGLOBETROTTER's admixture probability curves for Europe clusters 1-8. The cluster number, with sample size in parenthesis, and top two countries represented in the cluster are given in the top line. The black lines in each plot show the (scaled) probability, within that cluster's individuals, of matching two DNA segments to the reference populations listed just below the cluster name versus the centimorgan (cM) distance between the two DNA segments. Green lines depict the model fit when assuming a single pulse (date) of admixture, cyan lines when assuming a single pulse of admixture between only two sources, and red lines when assuming two pulses of admixture with distinct dates. There are two plots per cluster. The left plot depicts curves for the monotonically increasing reference pairing with highest r-squared between the green and black lines. The right plot depicts the same among reference pairings containing an African (plot title in bold) or East Asian (plot title in italics) population. In the left plot, we provide the inferred type of admixture and date(s) (B=BCE, otherwise CE) with 95% CI.

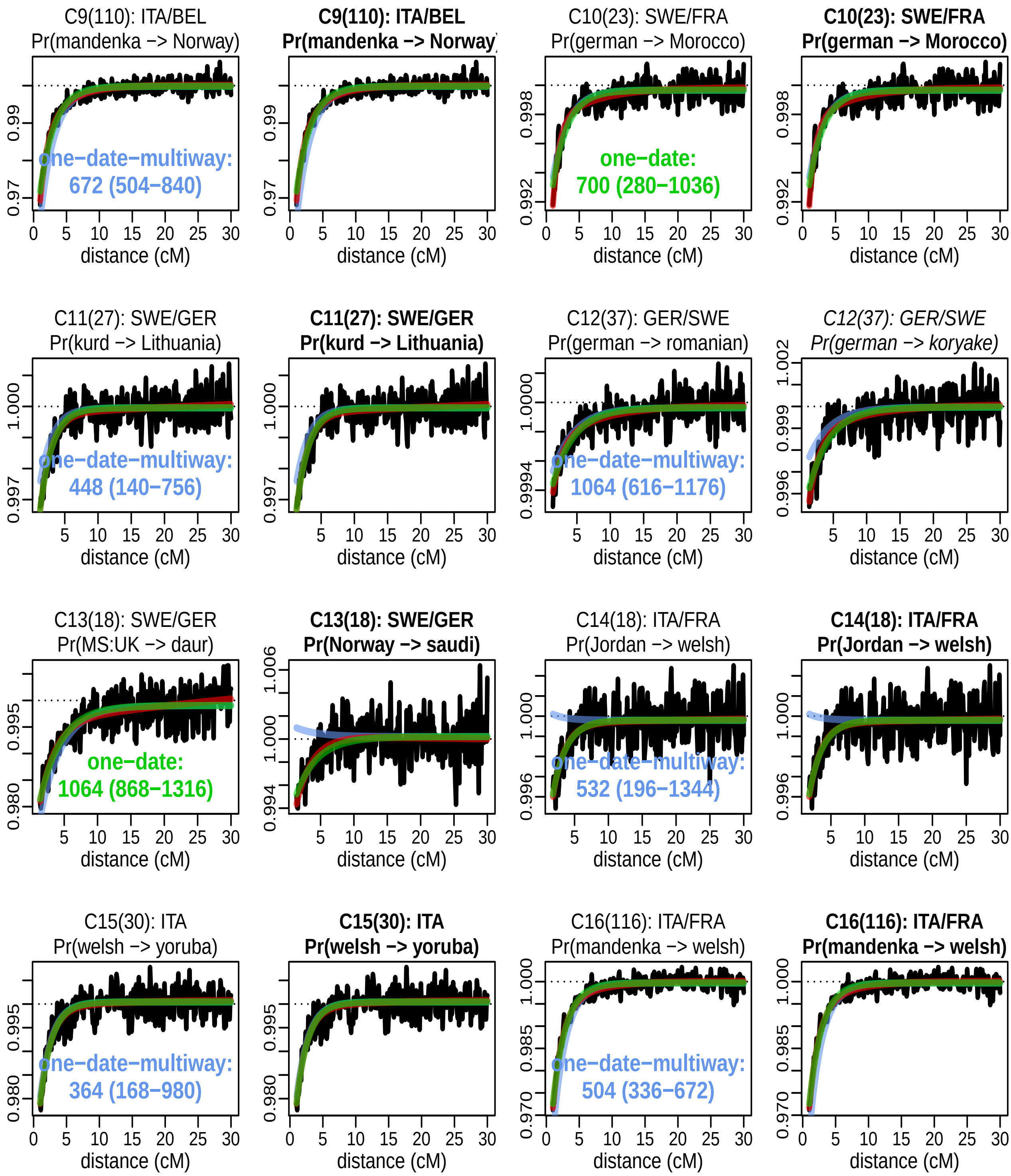


Figure S7: FastGLOBETROTTER's admixture probability curves for Europe clusters 9-16. See Fig S6 legend for details.

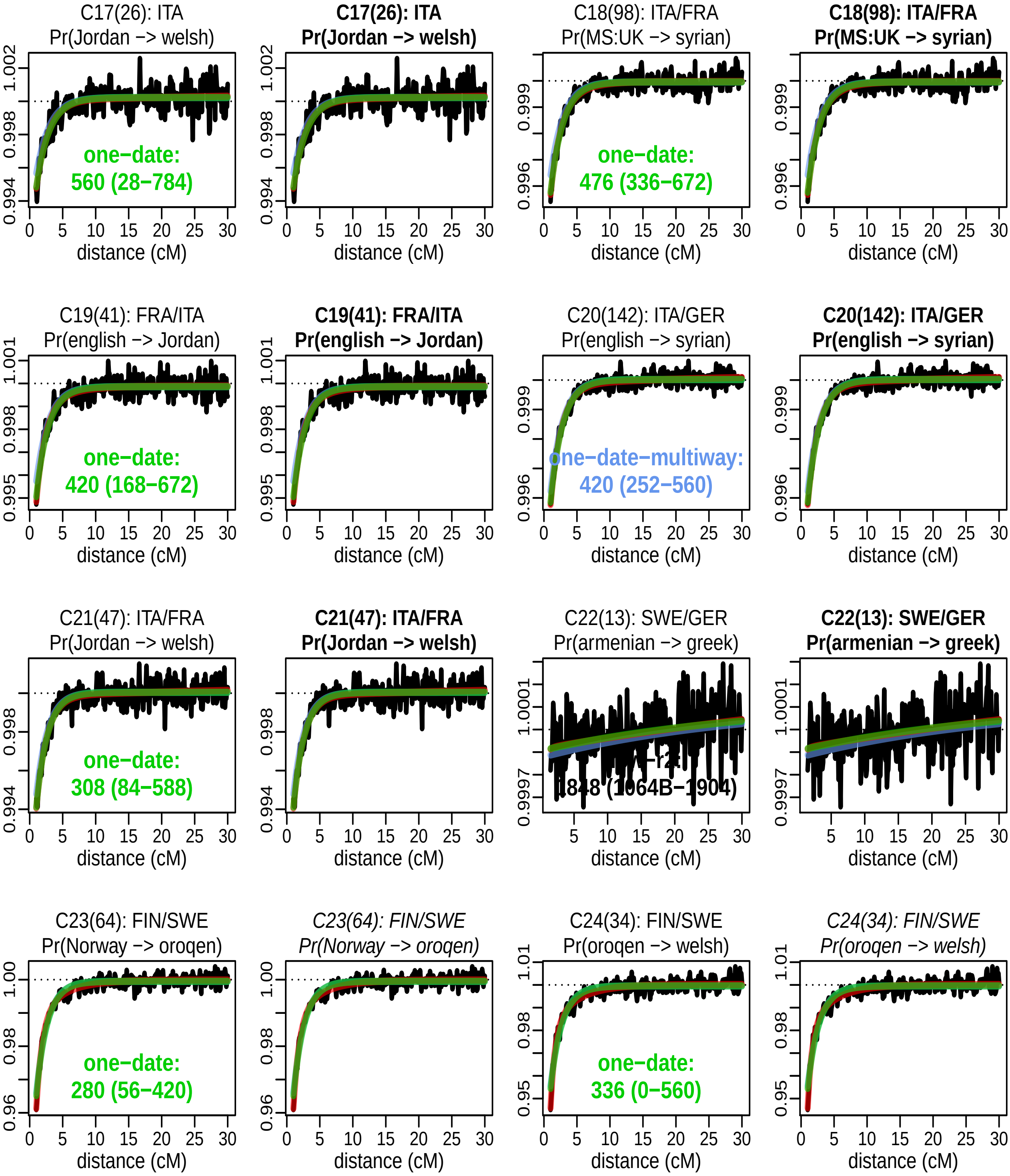


Figure S8: FastGLOBETROTTER's admixture probability curves for Europe clusters 17-24. See Fig S6 legend for details.

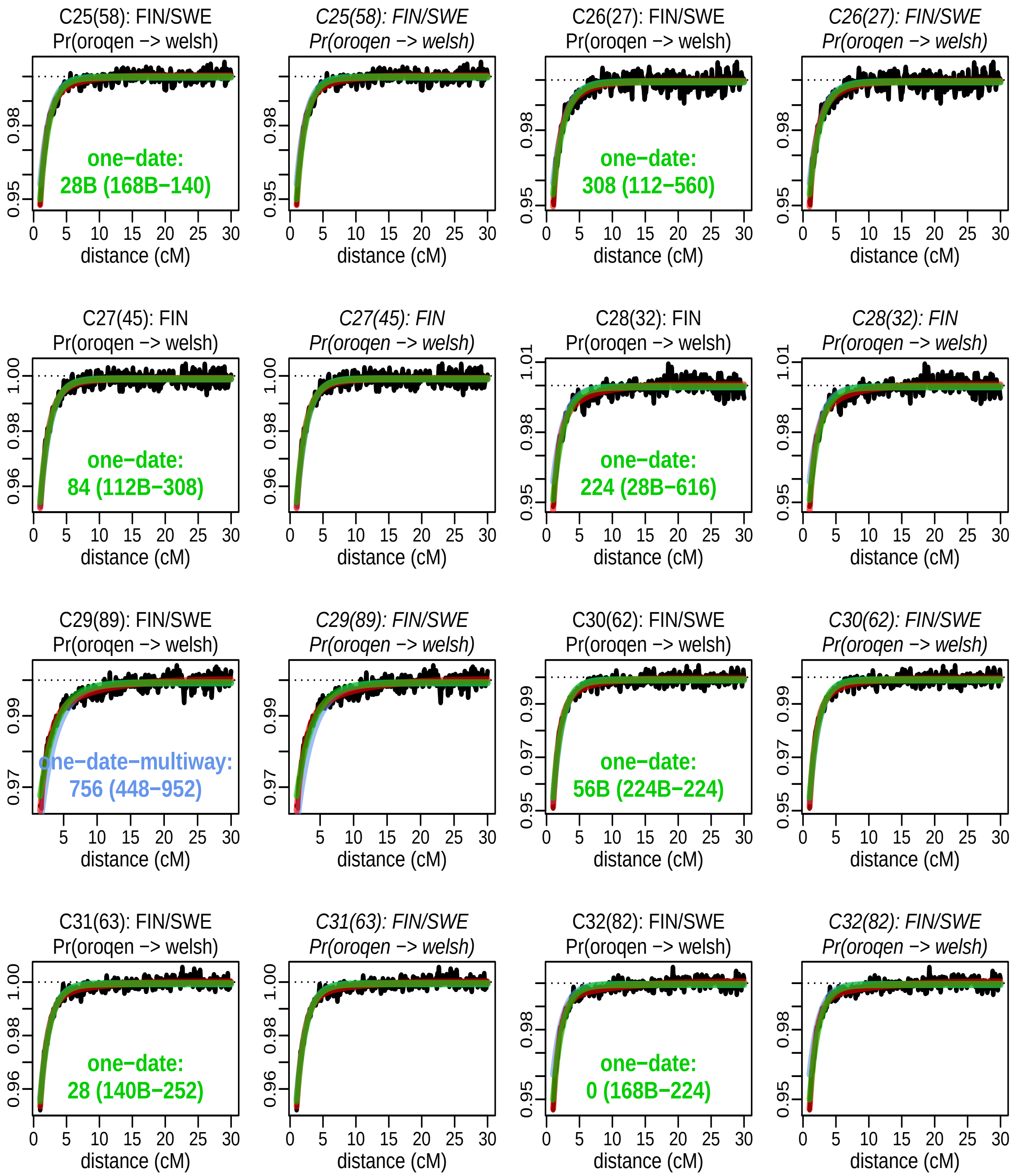


Figure S9: FastGLOBETROTTER's admixture probability curves for Europe clusters 25-32. See Fig S6 legend for details.

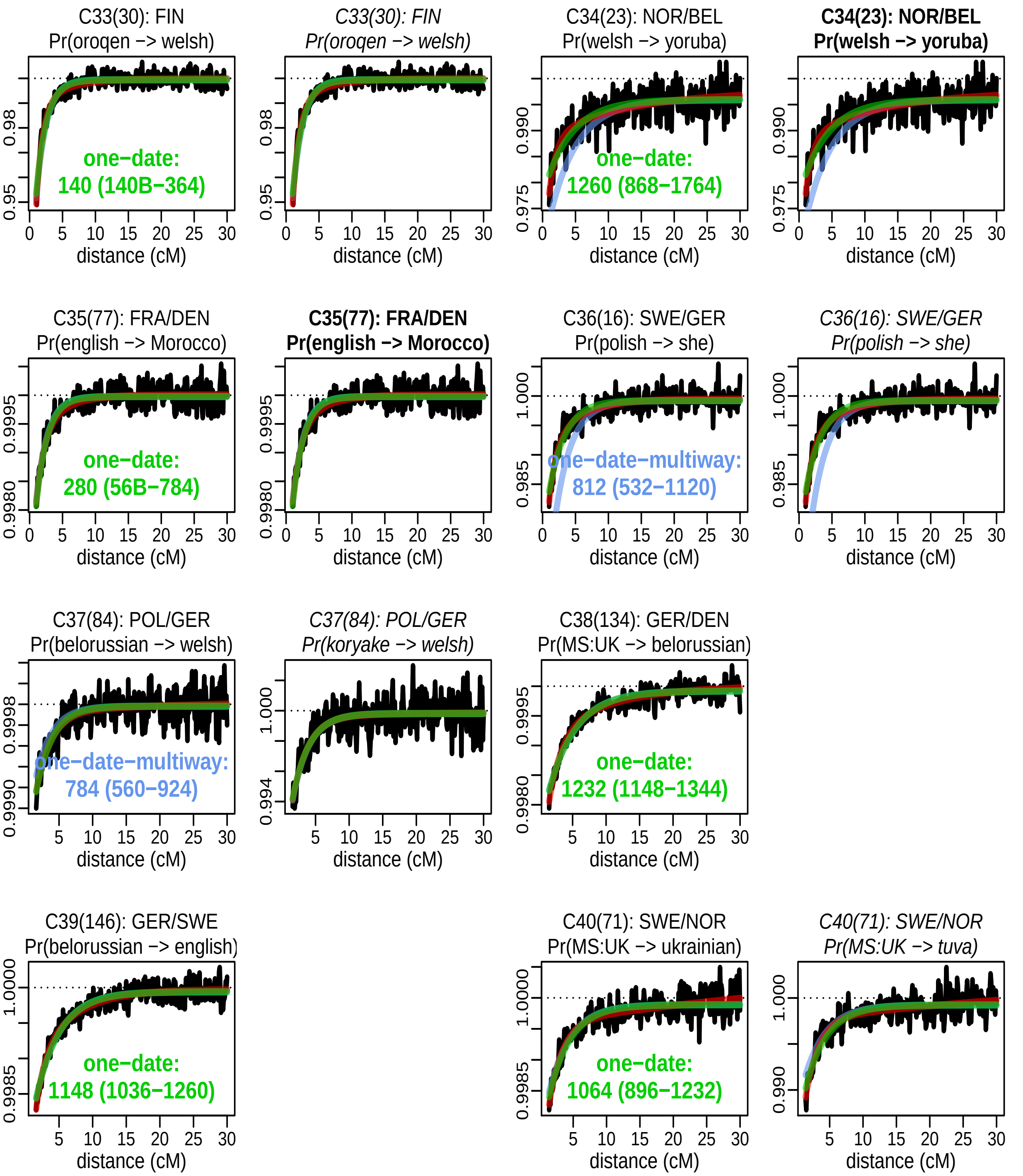


Figure S10: FastGLOBETROTTER's admixture probability curves for Europe clusters 33-40. See Fig S6 legend for details.

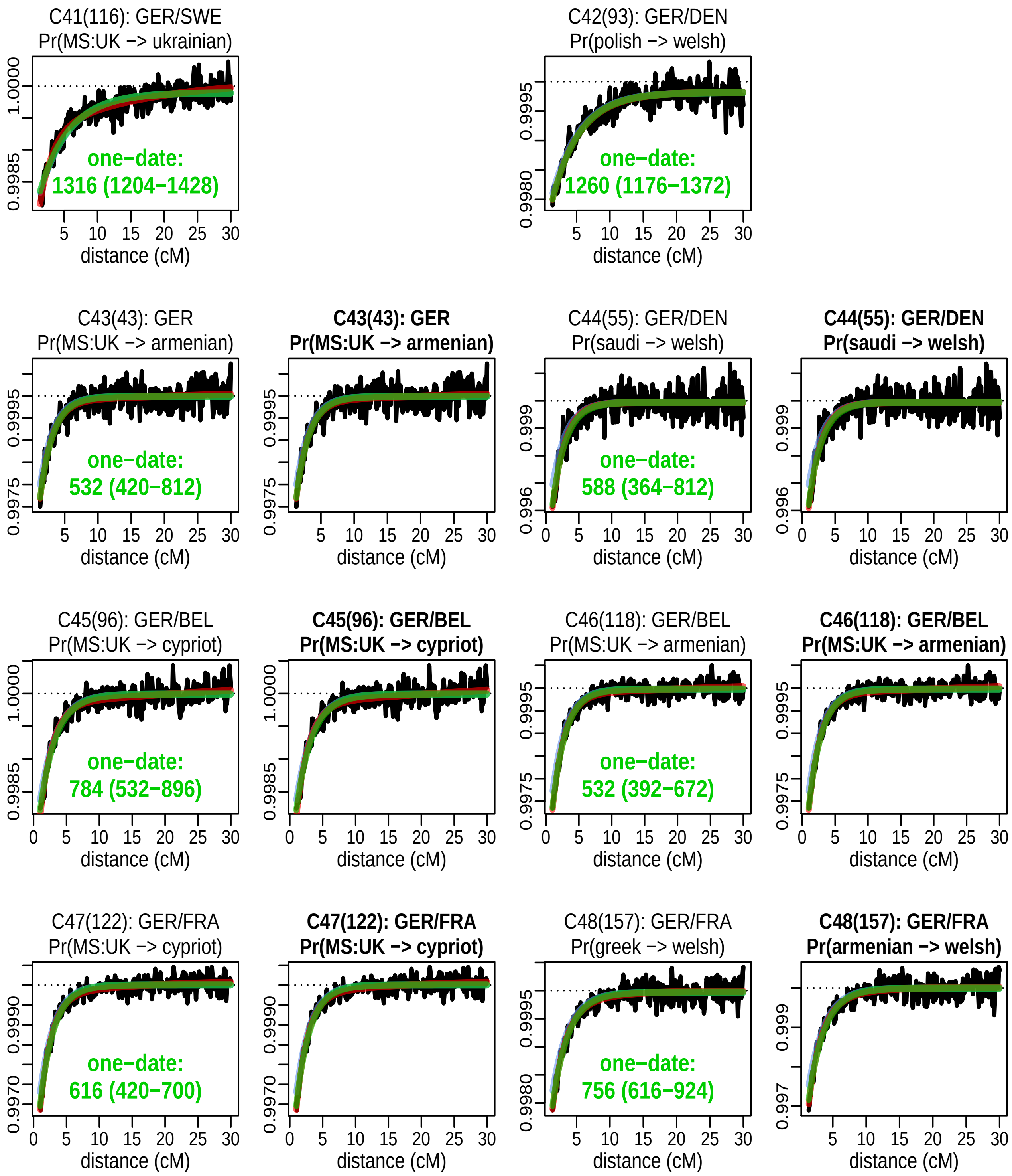


Figure S11: FastGLOBETROTTER's admixture probability curves for Europe clusters 41-48. See Fig S6 legend for details.

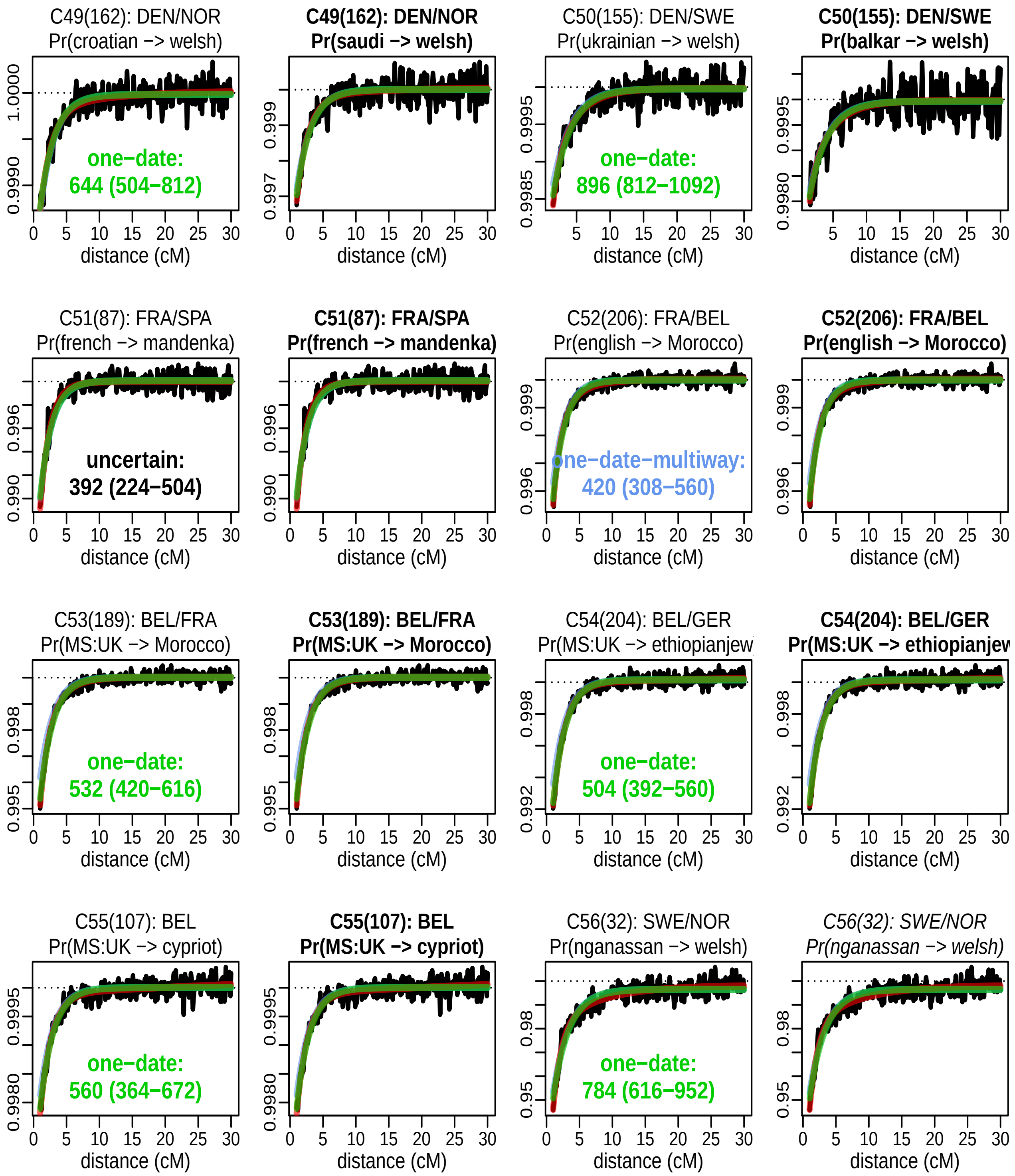


Figure S12: FastGLOBETROTTER's admixture probability curves for Europe clusters 49-56. See Fig S6 legend for details.

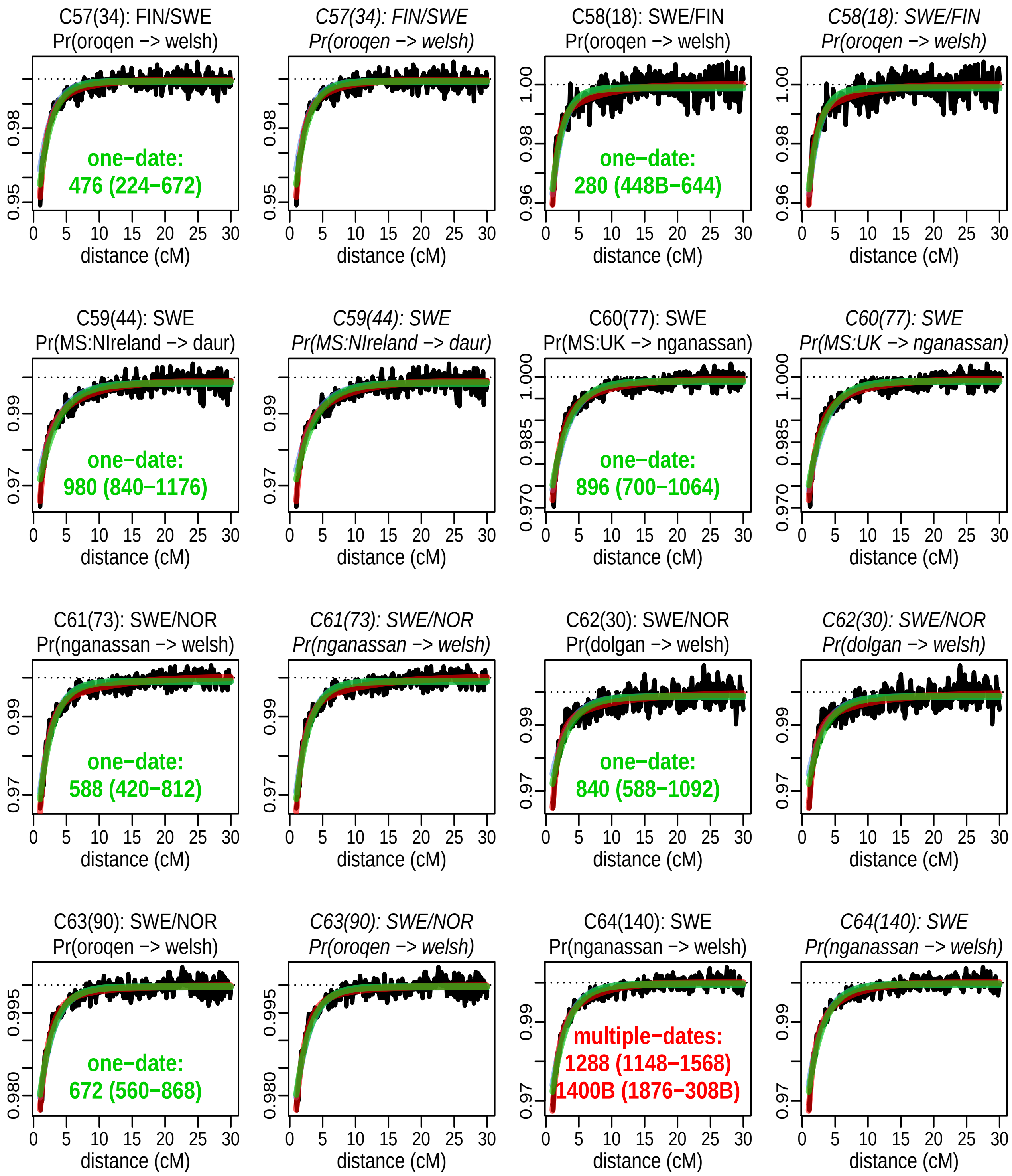


Figure S13: FastGLOBETROTTER's admixture probability curves for Europe clusters 57-64. See Fig S6 legend for details.

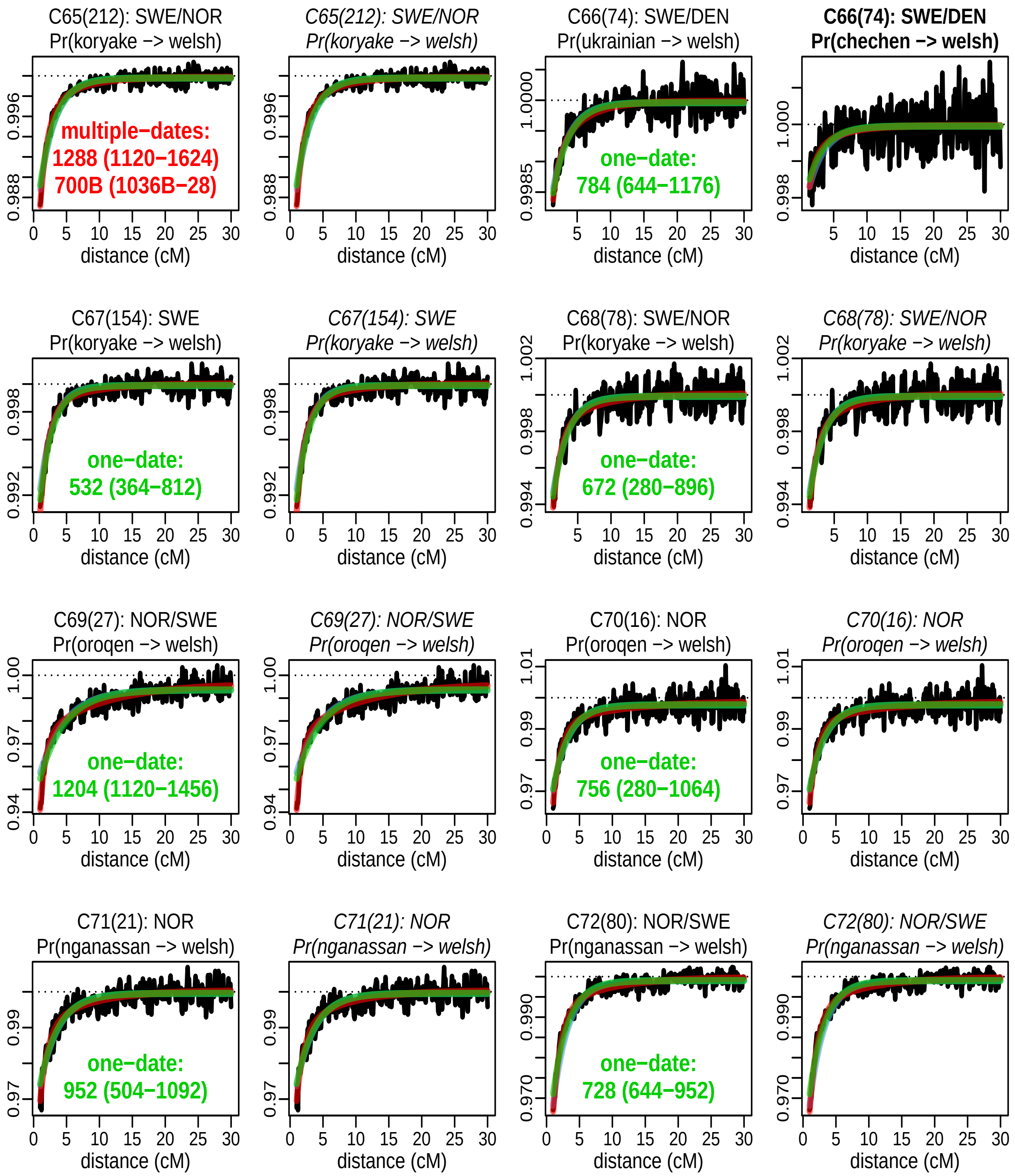


Figure S14: FastGLOBETROTTER's admixture probability curves for Europe clusters 65-72. See Fig S6 legend for details.

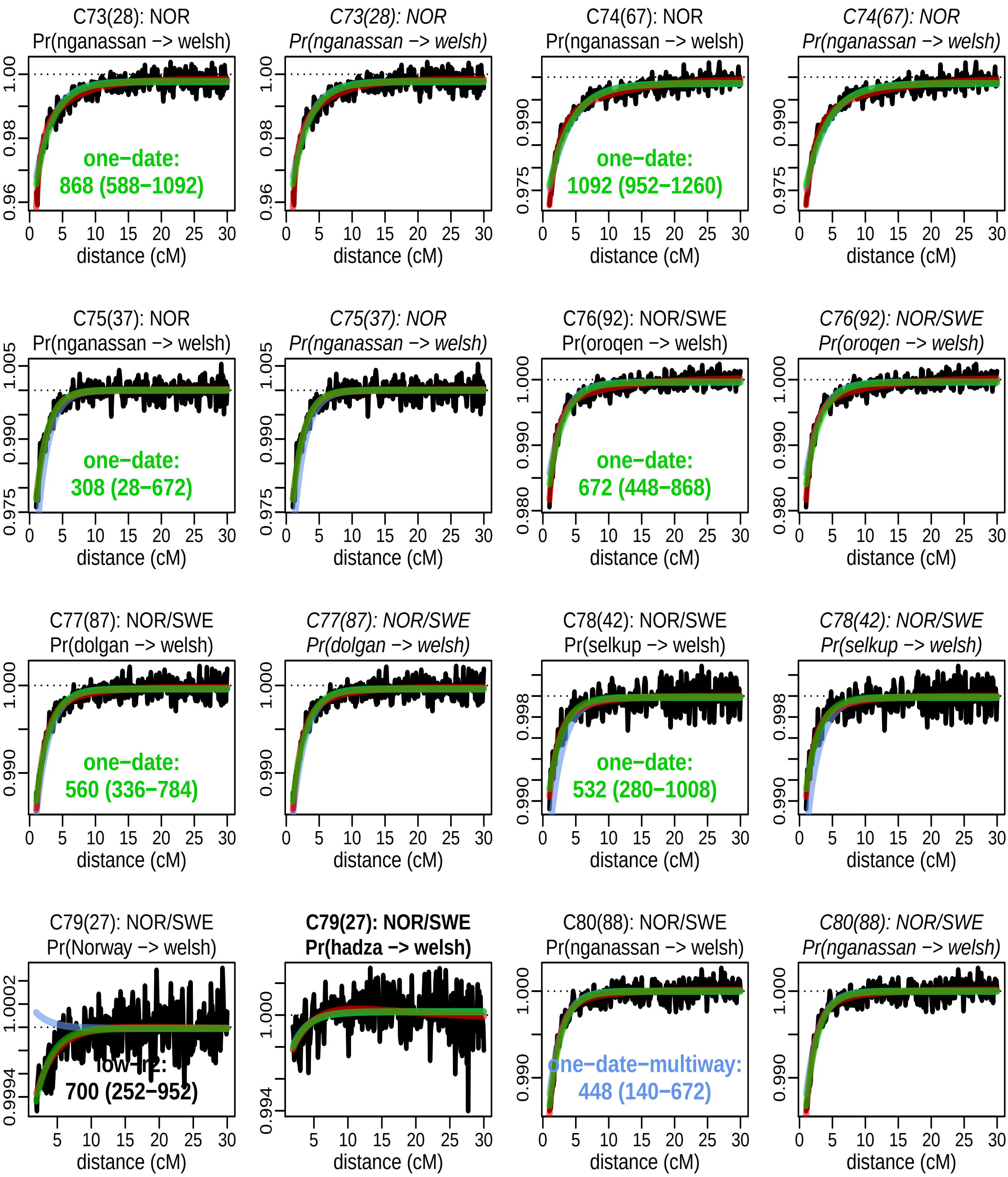


Figure S15: FastGLOBETROTTER's admixture probability curves for Europe clusters 73-80. See Fig S6 legend for details.

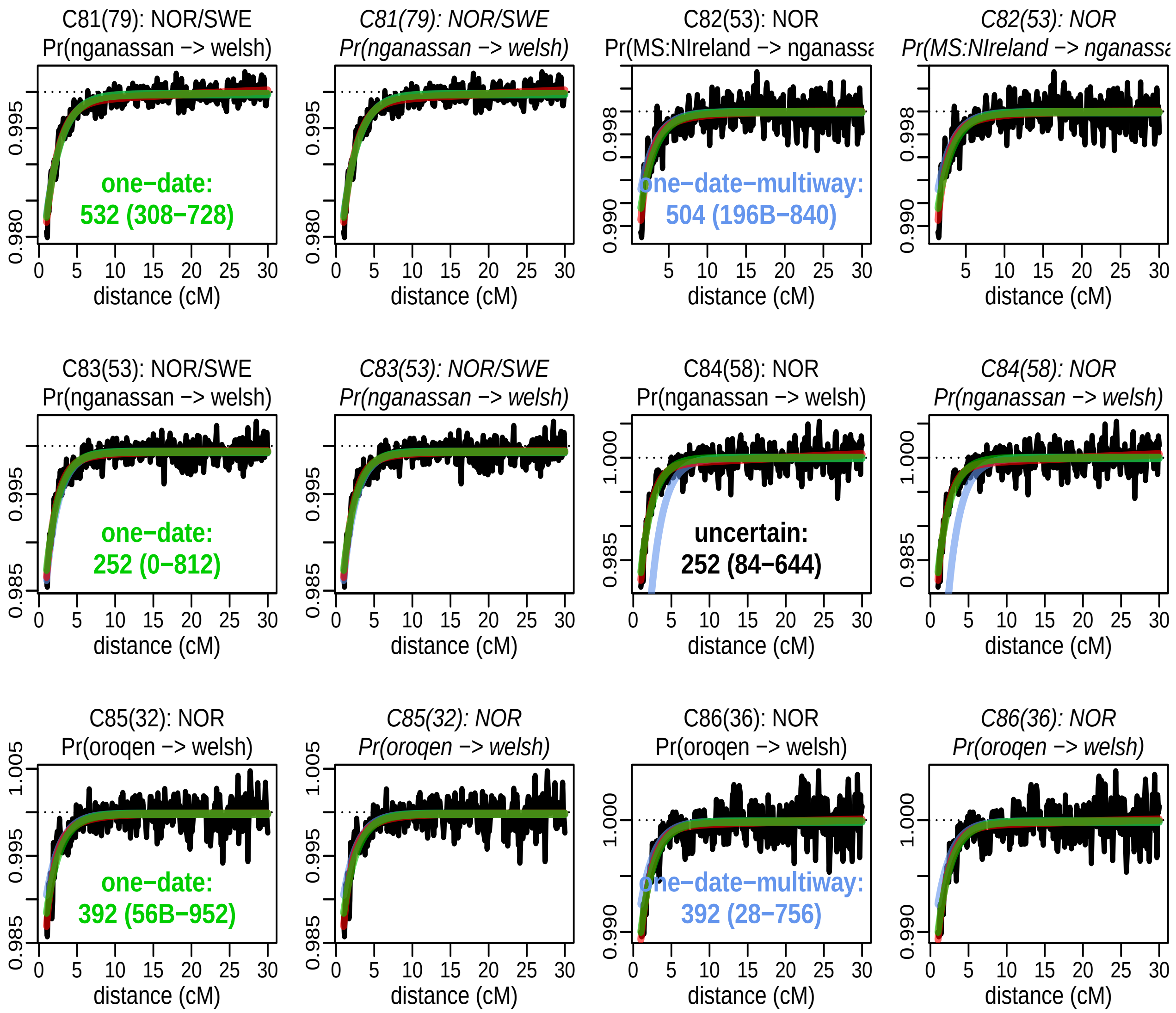


Figure S16: FastGLOBETROTTER's admixture probability curves for Europe clusters 81-86. See Fig S6 legend for details.
